# Cooperative phagocytosis underlies macrophage immunotherapy of solid tumours and initiates a broad anti-tumour IgG response

**DOI:** 10.1101/2022.01.01.474150

**Authors:** Jason C. Andrechak, Lawrence J. Dooling, Brandon H. Hayes, Siddhant Kadu, William Zhang, Ruby Pan, Manasvita Vashisth, Jerome Irianto, Cory M. Alvey, Dennis E. Discher

## Abstract

Macrophages are abundant in solid tumours and typically associate with poor prognosis, but macrophage clusters in tumour nests have also been reported as beneficial even though dispersed macrophages would have more contacts with cancer cells. Here, by maximizing both phagocytic activity and macrophage numbers, we discover cooperative phagocytosis by low entropy clusters in rapidly growing engineered immuno-tumouroids. The results fit the calculus of proliferation-versus-engulfment, and rheological measurements and molecular perturbations provide a basis for understanding phagocytic disruption of a tumour’s cohesive forces in soft cellular phases. The perturbations underscore the utility of suppressing a macrophage checkpoint in combination with an otherwise ineffective tumour-opsonizing monoclonal antibody, and the approach translates *in vivo* to tumour elimination that durably protects mice from re-challenge and metastasis. Adoptive transfer of engineered macrophages increases the fraction of mice that eliminate tumours and potentially overcomes checkpoint blockade challenges in solid tumours like insufficient permeation of blocking antibodies and on-target, off-tumour binding. Finally, anti-cancer IgG induced *in vivo* are tumour-specific but multi-epitope and contribute to a phagocytic feedback that drives macrophage clustering *in vitro*. Given that solid tumours remain challenging for immunotherapies, durable anti-tumour responses here illustrate unexpected advantages in maximizing net phagocytic activity.

## Introduction

The ability of a macrophage to engulf another cell or microbe in a tissue maintains homeostasis and potentially provides a first line of immune defense^1, 2^. However, within a solid tumour, if a macrophage is to physically engulf a cancer cell, then phagocytic forces^3^ must exceed the mechanical strength of the cohesion between solid tumour cells^4^. A large imbalance of such cell-cell interactions in mixtures of diverse tissue cell systems has long been seen to drive phase separation^5^, but segregation of immune cells is understudied. Clusters of macrophages in tumour nests have been reported to correlate with patient survival for at least two solid tumour types^6, 7^, and macrophage aggregation in the contexts of tissue injury has been compared to platelet clots^8^ that suggests a collective mechanical mechanism. On the other hand, tumour-associated macrophages more typically correlate with poor clinical prognoses^9^, and such macrophages not only promote growth and invasion in some cancers^10^ but also often lack phagocytic function^11^.

Phagocytosis of ‘self’ cells is generally inhibited by a key macrophage checkpoint interaction between SIRPα on the macrophage and CD47 on all cells including cancer cells^11–13^. Tumour cell engulfment can nonetheless be driven by anti-tumour monoclonal antibodies that bind Fc-receptors on macrophages^14^ (e.g. anti-CD20 in lymphoma). Importantly, more of these patients benefit when the monoclonal antibody is combined with antibody-based blockade of CD47^15^. However, response rates with similar combinations in solid tumours are significantly lower^16, 17^. Many phagocytosis signaling pathways are well-studied for macrophages attached to a culture dish^18–20^, but understanding of biophysical factors that influence phagocytosis in tumours is lacking. We hypothesized that a high number of maximally phagocytic macrophages could somehow work together – perhaps as clusters – to overcome both the cohesive forces and the proliferation of solid tumours.

Further challenges for macrophage checkpoint blockade in solid tumours include low permeation of anti-CD47^21^ relative to the potency of inhibitory signaling^22, 23^ as well as on-target, off-tumour binding of antibodies to ubiquitously expressed CD47. Adoptive transfer of engineered macrophages that traffic to tumours could potentially overcome both challenges to improve efficacy and safety of checkpoint blockade, especially if dosing maximizes the number of phagocytic macrophages and any cooperative effects. Unknown is whether these cells that constitute a first line of immune defense *in vivo* but lack antigen specific immune receptors could also contribute to any form of anti-cancer immunity^24, 25^. Given the requirement for combination of checkpoint disruption with tumour-opsonizing IgG monoclonal antibodies, we hypothesized *de novo* generation of anti-cancer IgG could be an effective form of acquired immunity. Our results provide the first evidence for *cooperative phagocytosis* by macrophages engulfing solid tumour targets and for an immune memory from macrophage checkpoint blockade that includes cancer-opsonizing IgG’s that drive macrophage clustering and tumour cell engulfment.

## Results

### Macrophages cluster when eradicating tumouroids

To determine the requirements for macrophages to eliminate a proliferating, cohesive mass of cancer cells, we engineered ‘tumouroids’ of B16 mouse melanoma cells in non-adhesive culture plates (**Fig. 1A**). This widely used tumour model for cancer immunotherapy development does not respond *in vivo* to T cell checkpoint blockade nor to CD47 disruption^26^ (and therefore represents a large number of patients thus far unresponsive to immunotherapy^27^). B16 cells adhere to each other and grow as thin dark tumouroids with irregular borders (Extended Data Fig. S1A), akin to early-stage melanoma in the upper layer of human skin^28^. Tumouroids resist deformation as a single cohesive mass when pulled into a micropipette (**Fig. 1B**), with a softness consistent with other neuro-lineage tissues such as brain that also relies for cohesion more on cell-cell adhesion than matrix adhesion.^29^ Sustainable stresses of ∼0.5-1 kPa are much higher than those exerted by a macrophage engulfing a microparticle^3^, but tumouroids flow after a few minutes as the stress favors disruption of adhesions – thus illustrating a means by which added macrophages can over time extract B16s from the tumouroid. Such viscoelastic behavior is typical of culture-aggregated spheroids, including cell mixtures that sort based on differential adhesion and cortical tension^30, 31^. Indeed, disrupting either Ca^2+^-dependent cell adhesions or actin polymerization relaxed tumouroids as evidenced by a rapid (<1 h) increase in projected area (Extended Data Fig. 1B). Subcutaneous tumours of B16s from mice exhibit similar softness (Extended Data Fig. 1C-F), despite the presence of various other cell types including macrophages.

**Figure 1:**
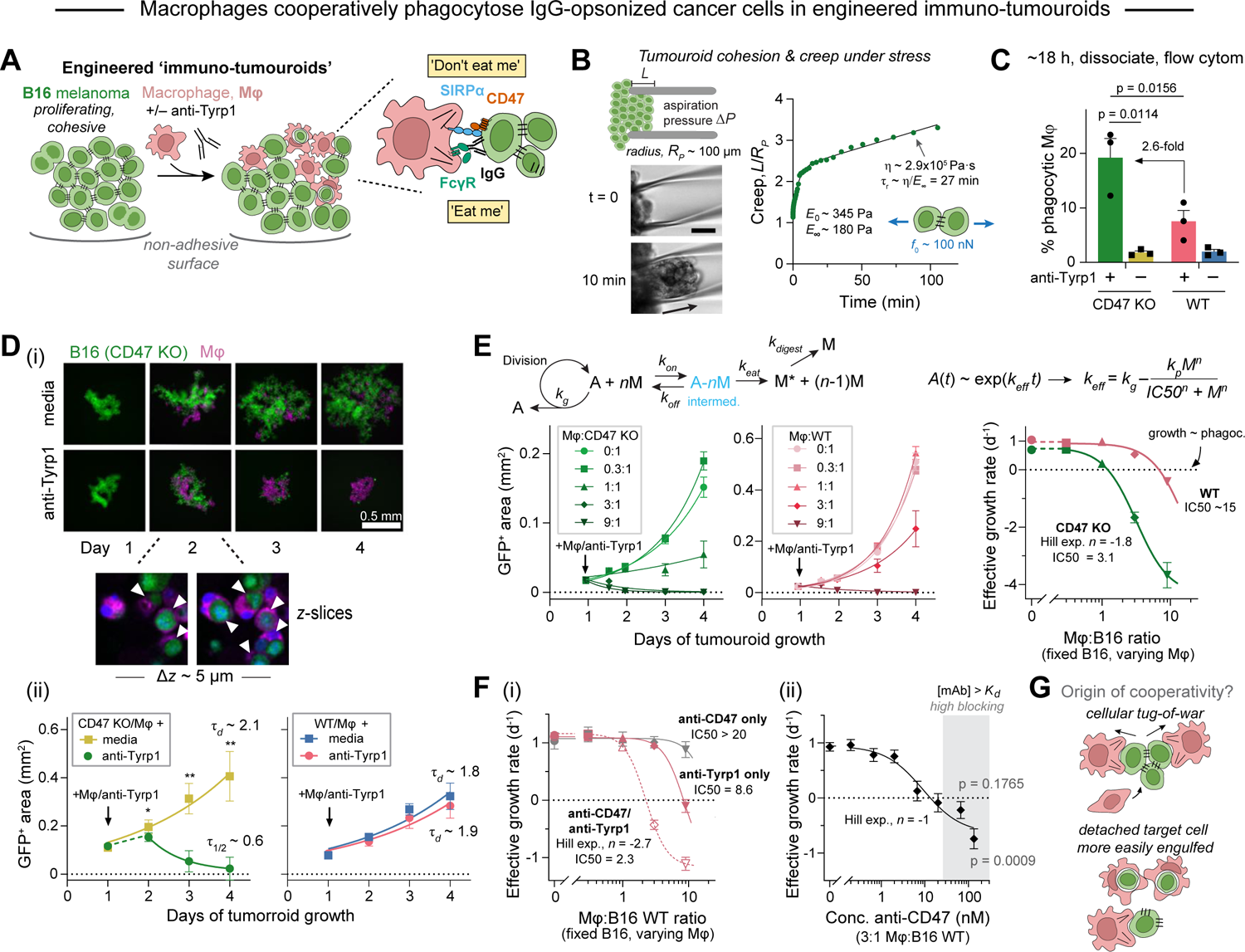
Macrophages cooperatively phagocytose IgG-opsonized cancer cells in engineered tumouroids. **A** Engineered ‘immuno-tumouroid’ model for phagocytosis of cohesive tumour cells. Tumouroids were formed by culturing B16 melanoma cells on non-adhesive surfaces and subsequent addition of bone marrow-derived macrophages with or without opsonizing anti-Tyrp1 IgG. The inset depicts a macrophage-melanoma tumouroid ‘phagocytic synapse’ with FcR-IgG signaling that promotes phagocytosis and CD47:SIRPα that inhibits phagocytosis. **B** Micropipette aspiration of B16 tumouroids at *t* = 0 and *t* = 10 min of constant stress (aspiration pressure, Δ*P)*. Tumouroid strain or creep is plotted vs. time and fit with a standard linear model to determine the elastic moduli and viscosity. Scale bar: 50 µm. Forces on the order of 100 nN are predicted to be sustained by tumouroids without rupture based on typical aspiration pressures. **C** Phagocytosis of CD47 KO and WT B16 tumouroids ∼18 h after addition of macrophages at a nominal 1:1 macrophage:B16 ratio with or without anti-Tyrp1. The % phagocytic macrophages corresponds to GFP+ macrophages in cell suspensions from disaggregated tumouroids (mean ± SD, n = 3 where each replicate consists of cells pooled from the same 96-well plate). Statistical significance was assessed by two-way ANOVA and Tukey’s multiple comparison test. **D** Opsonization with anti-Tyrp1 limits the growth of CD47 KO ‘immuno-tumouroids’, but not WT immuno-tumouroids. **(i)** Representative fluorescence images depict growth or repression of CD47 KO (green) in immuno-tumouroids from days 1-4 when untreated (top row) or treated with anti-Tyrp1 (bottom row). Macrophages (magenta, ∼1:1 ratio to initial B16 number) and anti-Tyrp1 were added immediately after the day 1 images were acquired. Scale bar: 0.5 mm. Confocal z-slices of CD47 immuno-tumouroids on day 2 clearly depict B16s engulfed by macrophages (white arrows indicate engulfed cell). **(ii)** Tumouroid growth was measured by calculating the projected GFP+ area at indicated time points (mean ± SD, n = 6 tumouroids). Statistical significance was assessed by the Mann-Whitney test (unpaired, two-tailed, * p = 0.03, ** p = 0.0043) comparing GFP+ areas with and without anti-Tyrp1 at each time point. Solid lines are nonlinear regression of the data to a simple exponential of the form *A*(*t*) = *A*_1_ exp[*k(t*-1)]. The doubling time *τ_d_* or half-life *τ*_1/2_ was calculated from the fitted rate constant *k*. **E** A reaction-kinetic model accounts for division of cancer cells (*A*) in tumouroids and phagocytosis by n macrophages (*M*). Exponential growth or decay of tumouroid area (fitted curves) depends on the ratio of macrophages to B16 cells as well as CD47 on target cells (left) (mean ± SEM, n = 6-8 tumouroids from a representative of four or more experiments). The effective growth rate (*k_eff_*) from fitted exponentials of CD47 KO tumouroids exhibits a Hill-like dependence on macrophage number (Hill parameter = −1.8, IC50 = 3.1, mean ± SEM, n ≥ 24 tumouroids from four independent experiments). A constrained fit of WT tumouroid growth with the same Hill parameter indicates much higher macrophage:B16 ratios are required for tumouroid elimination (IC50 = ∼15) (right). **F (i)** Effective growth rates of WT tumouroids with anti-CD47 and anti-Tyrp1 also show Hill-like dependence on macrophage number (Hill parameter = −2.7, IC50 ratio = 2.3, mean ± SEM, n ≥ 18 tumouroids across three independent experiments). The concentration of anti-CD47 (133 nM) used is approximately saturating for CD47 on WT B16 as shown in Fig S2A-ii. **(ii)** The effective growth rate of WT tumouroids at a fixed 3:1 macrophage:B16 ratio depends on the concentration of anti-CD47, but tumouroid elimination is not cooperative with respect to blocking Ab level (mean ± SEM, n ≥ 16 tumouroids across three independent experiments). Statistically significant growth repression (*k_eff_* < 0, one-sample t-test) is only observed at the highest level of blocking (>100 nM anti-CD47 as used in Fig. F-i) that exceeds by ∼4-fold the measured affinity of blocking Ab for CD47 (*K* ∼ 25 nM as shown in Fig. S2A-ii). **G** Proposed cooperative tug-of-war by maximally phagocytic macrophages to **(i)** disrupt target cancer cell adhesions and subsequently **(ii)** phagocytose individual cells.

We hypothesized that cancer phagocytosis could be maximized for targets lacking CD47, and we therefore generated tumouroids from CRISPR/Cas9-engineered B16 cells with either *Cd47* knockout (KO) or wild-type (WT) levels of CD47 (Extended Data Fig. 2A). Bone marrow-derived macrophages were added to pre-assembled tumouroids together with anti-Tyrp1 monoclonal antibody that binds and opsonizes B16 cells (Extended Fig. 2A-B). Tyrp1 aids in melanin granule synthesis, so that anti-Tyrp1 is relatively specific to melanocytes in contrast to the ubiquitous phagocytosis checkpoint ligand CD47 (Extended Data Fig. 2C). Despite the specificity, anti-Tyrp1 was recently found ineffective in the clinic against melanoma^32^ and shows little to no effect on B16 tumours established in mice^26^. Growth of WT tumouroids is likewise unaffected by anti-Tyrp1 with or without added macrophages, *but* we do find opsonization of CD47 KO ‘immuno-tumouroids’ maximizes phagocytic macrophages quantified ∼1-day after addition (**Fig 1C**) (Extended Data Fig. 2D). Conventional assays with cancer cell suspensions added to immobilized macrophages for 1-2 h show similar phagocytosis trends (Extended Data Fig. 2E) – all of which suggests macrophages in tumouroids can extract and engulf individual B16s, as clearly visualized for opsonized CD47 KO tumouroids (**Fig 1D-i**). Importantly, only for this maximal phagocytosis condition is the exponential growth of the tumouroid reversed and tumouroids eradicated (**Fig 1D-ii**) (Extended Data Fig. 3A).

**Figure 2:**
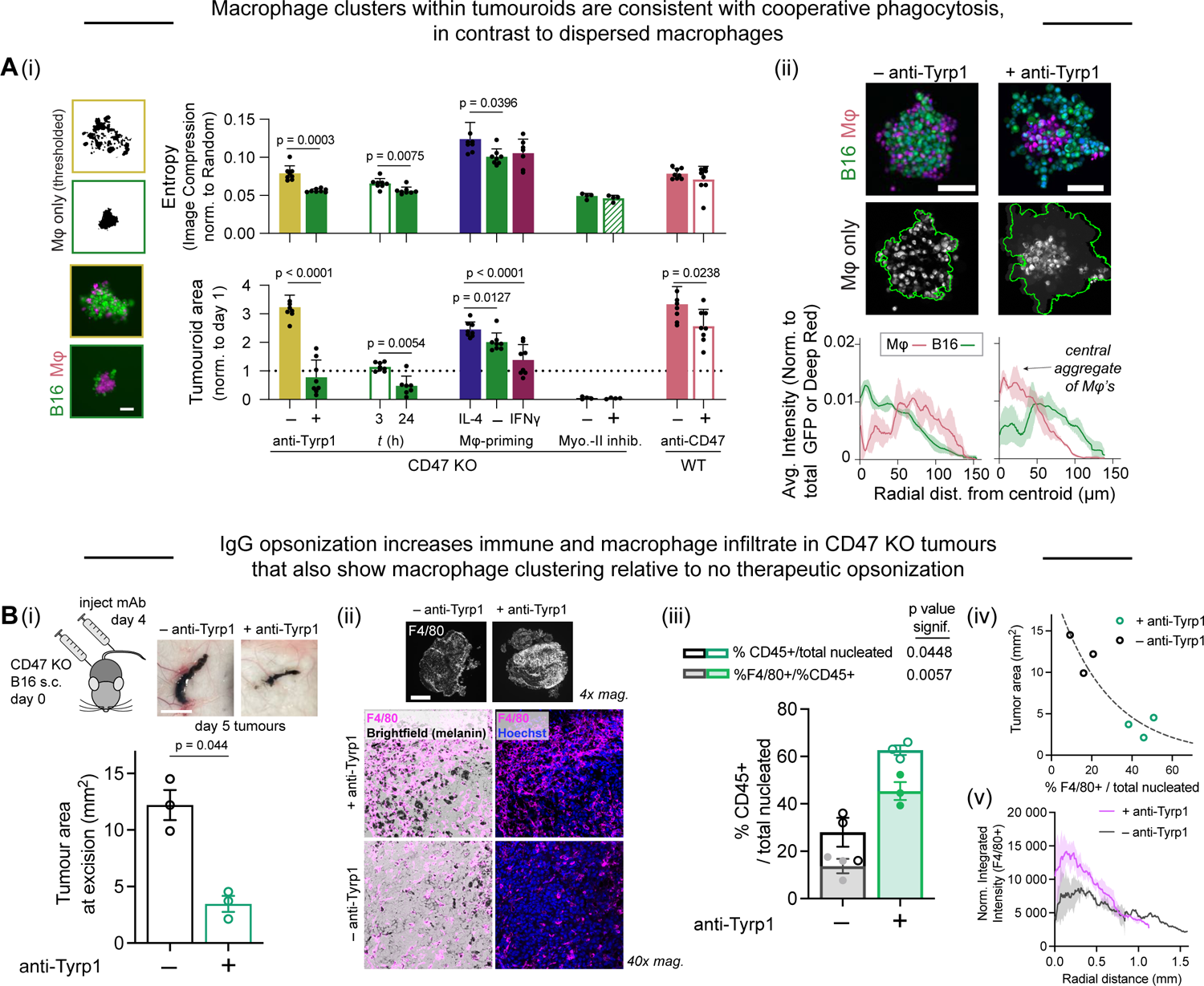
Macrophages infiltrate, cluster, and repress CD47-depleted syngeneic tumours *in vivo* only in combination with therapeutic opsonization by IgG. **A** Macrophage clusters in tumouroids are consistent with cooperative phagocytosis. **(i)** The informational entropy approximated by image file compression analysis (Extended Data Fig. 4) summarizes macrophage clustering within CD47 KO tumouroids across multiple experimental conditions including anti-Tyrp1 opsonization, kinetic studies, macrophage-priming with interferon-γ (IFNγ) or interleukin-4 (IL-4), and myosin-II inhibition with blebbistatin, as well as CD47 antibody blockade on WT tumouroids. Corresponding GFP+ tumouroid areas (normalized to areas on day 1) are shown for the same conditions (mean ± SD, n = 6-8 tumouroids). Statistical significance of entropy and tumour area differences were assessed by Welch’s t-test (two-tailed, unpaired) for comparisons between two experimental conditions and one-way ANOVA with Tukey’s multiple comparison test for the macrophage-priming experiment with three conditions. The representative thresholded macrophage images (top left) and fluorescence images of B16 and macrophage (bottom left) are from CD47 KO tumouroids ∼24 h after addition of 3:1 macrophage:B16 without anti-Tyrp1 (yellow frames) and with anti-Tyrp1 (green frames), which correspond to the first two bars in the graph. Scale bar: 100 µm. **(ii)** Representative max intensity projections of confocal fluorescence images (top) of CD47 KO (green) and macrophage (magenta) tumouroids one day after addition of macrophages with and without anti-Tyrp1. Scale bar: 100 µm. Radial profiles (bottom) of macrophage and B16 fluorescence within CD47 KO tumouroids indicate macrophage clustering occurs only when tumouroids are opsonized with anti-Typr1 (mean ± SD, n = 3). **B (i)** Representative photographs of untreated and anti-Tyrp1-treated tumours prior to disaggregation and analysis by flow cytometry (top). Scale bar: 1 mm. Tumour area as measured on untreated or anti-Tyrp1-treated mice on day 5 post inoculation, 24 h after treatment (bottom). Statistical significance was assessed by unpaired t-test (mean ± SEM, n = 3 per group). **(ii)** Representative fluorescence images of F4/80 staining in untreated or anti-Tyrp1-treated CD47 KO tumours on day 5, 24 h after treatment. Scale bar: 0.5 mm **(iii)** CD45 and F4/80 staining of isolated tumours 24 h after a single dose of anti-Tyrp1 on day 4 after tumour engraftment. Anti-Tyrp1-treatment tumours showed increased infiltrate of CD45+ cells as a fraction of total nucleated cells relative to untreated control, of which a ∼2-fold larger fraction of CD45+ cells were also F4/80+. Statistical significance was assessed by unpaired t-test with Welch’s correction (n = 3 per group for CD45+ staining) and by unpaired t-test (n = 3 per group for F4/80+ staining), mean ± SEM. **(iv)** Relationship between tumour area and F4/80+ macrophage immune infiltrate, fit with an exponential curve. F4/80+ cell frequency correlates negatively with tumour area on day 5. **(v)** Radial profile analysis of F4/80+ signal in CD47 KO tumours based on 4x images (n = 3 per group). F4/80+ cells infiltrate more highly into the core of anti-Tyrp1-treated tumours and more highly overall, while control tumours show evenly distributed, lower overall F4/80+ infiltrate.

**Figure 3:**
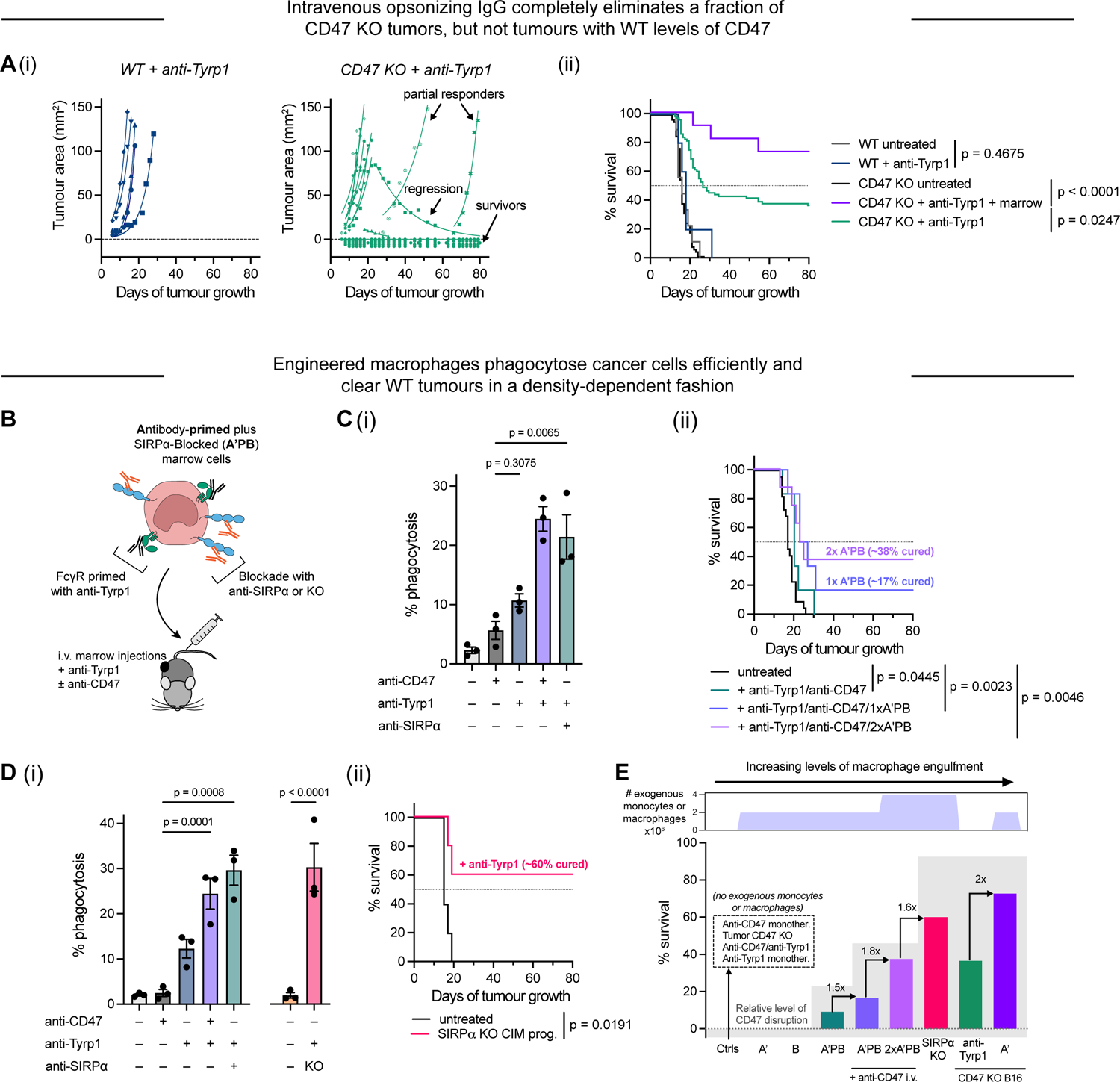
Increasing macrophages boosts complete anti-tumour response rates of CD47-depleted tumours; engineered macrophages phagocytose cancer cells efficiently and clear WT tumours in a density-dependent fashion. **A (i)** Representative tumour growth curves of projected tumour area versus days after tumour engraftment. Each symbol represents a separate tumour and is fit with an exponential growth equation A = A_0_^ekt^ (solid line). Complete anti-tumour responses in which a tumour was never palpable are all depicted with the same symbol (filled circle) and solid lines at A = 0. For KO + anti-Tyrp1, n = 15 across 2 independent experiments are shown; for B16 ctrl + anti-Tyrp1, n = 5. **(ii)** Survival curves of mice challenged with 2×10^5^ WT or CD47 KO cells subcutaneously (s.c.) and treated i.v. with 250 µg anti-Tyrp1 on days 4, 5, 7, 9, 11, 13, and 15 or left untreated. Anti-Tyrp1 treatment has no effect on WT tumour survival (treated n = 5, untreated n = 9), but can eliminate ∼40% of CD47 KO tumours (treated n = 82, untreated n = 60). Donor bone marrow injections combined with i.v. anti-Tyrp1 treatment (n = 11 across 2 independent experiments) increase the fraction of mice that show complete anti-tumour responses against CD47 KO tumours by 2-fold. Statistical significance was determined by the Log-rank (Mantel-Cox) test. **B** Scheme of antibody-primed (with anti-Tyrp1, A’), Plus (P) SIRPα-blocked (by anti-SIRPα, B) marrow cells (A’PB, left). Conditionally immortalized macrophage (CIM) progenitors were also used to generate SIRPα KO macrophage progenitors as an improvement of antibody-based blockade. Cells were then injected intravenously into tumour-bearing mice on day 4 (right) with indicated antibody i.v. injections on days 4, 5, 7, 9, 11, 13, and 15. **C (i)** Phagocytosis of WT B16s by bone marrow-derived macrophages. Macrophages with CD47-SIRPα signaling disrupted (through anti-CD47 or anti-SIRPα (A’PB)) and target IgG opsonization effectively engulf cancer cells *in vitro*. Statistical significance was assessed by one-way ANOVA with Sidak’s test for multiple comparisons (mean ± SEM, n = 3 per condition). **(ii)** Survival curves of WT tumour-bearing mice (non-KO guide ctrl B16) treated i.v. with anti-CD47 and anti-Tyrp1 combined with either a one (1x) or two (2x) doses of A’PB. The second A’PB dose was administered on day 7. Statistical significance was determined by the Log-rank (Mantel-Cox) test (untreated n = 22, anti-Tyrp1/anti-CD47 n = 6, anti-Tyrp1/anti-CD47/1x A’PB n = 6, anti-Tyrp1/anti-CD47/2x A’PB n = 8). **D (i)** Phagocytosis of WT B16s by CIMs. Unedited WT CIMs (*left)* engulf opsonized target B16s similar to bone marrow-derived macrophages with CD47-SIRPα disruption, while SIRPα KO CIMs also engulf target B16s highly with opsonization. Statistical significance was assessed by one-way ANOVA with Sidak’s test for multiple comparisons (mean ± SEM, n = 3 per condition). **(ii)** Survival curves of WT tumour-bearing mice (parental B16) treated i.v. with anti-Tyrp1 combined with a single dose of SIRPα KO CIM progenitors on day 4 with indicated antibody i.v. injections on days 4, 5, 7, 9, 11, 13, and 15. Significance was determined by the Log-rank (Mantel-Cox) test (n = 5 per group). **E** Summary of complete anti-tumour response data showing that when CD47 is disrupted to maximize macrophage engulfment, WT B16 tumours can be eradicated in a macrophage density-dependent fashion. High macrophage transfer counts and permanent ablation of CD47-SIRPα signaling in either engineered macrophage or in tumours results in the highest rate of tumour elimination.

### Macrophages cooperatively phagocytose targets

Surprisingly, macrophages segregate under maximal phagocytosis conditions (Fig 1D-i). Such a cluster will have few macrophages next to cancer cells and will physically impede interactions with cancer cells compared to the dispersed macrophages under the other conditions. Despite the macrophage clustering, adding more macrophages suppresses growth and fits the calculus of cell proliferation suppressed on a stoichiometric basis by *n* phagocytes (**Fig 1E**) (Extended Data Fig. 3B). The rate constant *k_eff_* exhibits a Hill slope *n =* −1.8 for KOs that indicates *cooperative* phagocytosis as a function of macrophage number (Extended Data Fig. 3C,D). WT tumouroids could be eliminated by high concentrations of anti-CD47 + anti-Tyrp1 plus high macrophage numbers, and tumouroid area again indicated cooperativity as a function of added macrophages (**Fig 1F-i**) but not as a function of anti-CD47 (**Fig 1F-ii** and Extended Data Fig. 3D-E). High dose anti-CD47 blockade can be difficult to achieve clinically given the low permeation of solid tumours^4^, and the results also underscore a cell-level cooperativity that reflects a cell-level structure such as a macrophage cluster. Indeed, although anti-CD47 + anti-Tyrp1 enhanced engulfment of WT B16 suspensions in standard 2D phagocytosis assays relative to controls, high-density regions of the immobilized macrophages showed *fewer* phagocytic macrophages than low-density regions – contrary to cooperativity (Extended Data Fig. 3F). Tumouroid cohesion might be needed for macrophage cooperativity, with macrophage clusters providing a structural basis for this functional cooperativity – in a hypothetical “tug-of-war” on cohesive tumouroids by macrophage before successful phagocytosis of the targets (**Fig 1G**).

To assess whether macrophage clustering consistently associates with tumouroid suppression, an information entropy approach^33^ was applied to our many images obtained under a wide range of time points and perturbations. Image files of clustered macrophages are digitally compressed more than images of dispersed macrophages (i.e. more disordered) (**Fig.2A-i** and Extended Data Fig. 4), and tumouroid area measurements at the same timepoint show the same trend. Cell density profiles further show a central cluster of macrophages displaces B16 cells in tumouroids only under maximum engulfment conditions (**Fig.2A-ii**), and kinetic studies indicate clustering takes more than 3 h but is clear by 24 h (Fig 2A-i). Tumouroid elimination and tight macrophage clusters are also promoted by pre-treating cultured macrophages with interferon-γ (IFNγ) (Fig 2A-i), which increases macrophage surface receptors and results in a low contractility mechanobiological state^11^ consistent with soft tumouroid engulfment (e.g. Myosin-II inhibition has no effect); interleukin-4 (IL-4) has the opposite effects and disperses macrophages (Extended Data Fig. 5). The macrophage priming approach notably avoids direct cytokine effects on B16 growth^34^. Further studies show macrophage clusters are not sufficient for efficient phagocytosis and tumouroid elimination, but the various analyses do indicate that: when macrophage engulf maximally, they cluster.

**Figure 4:**
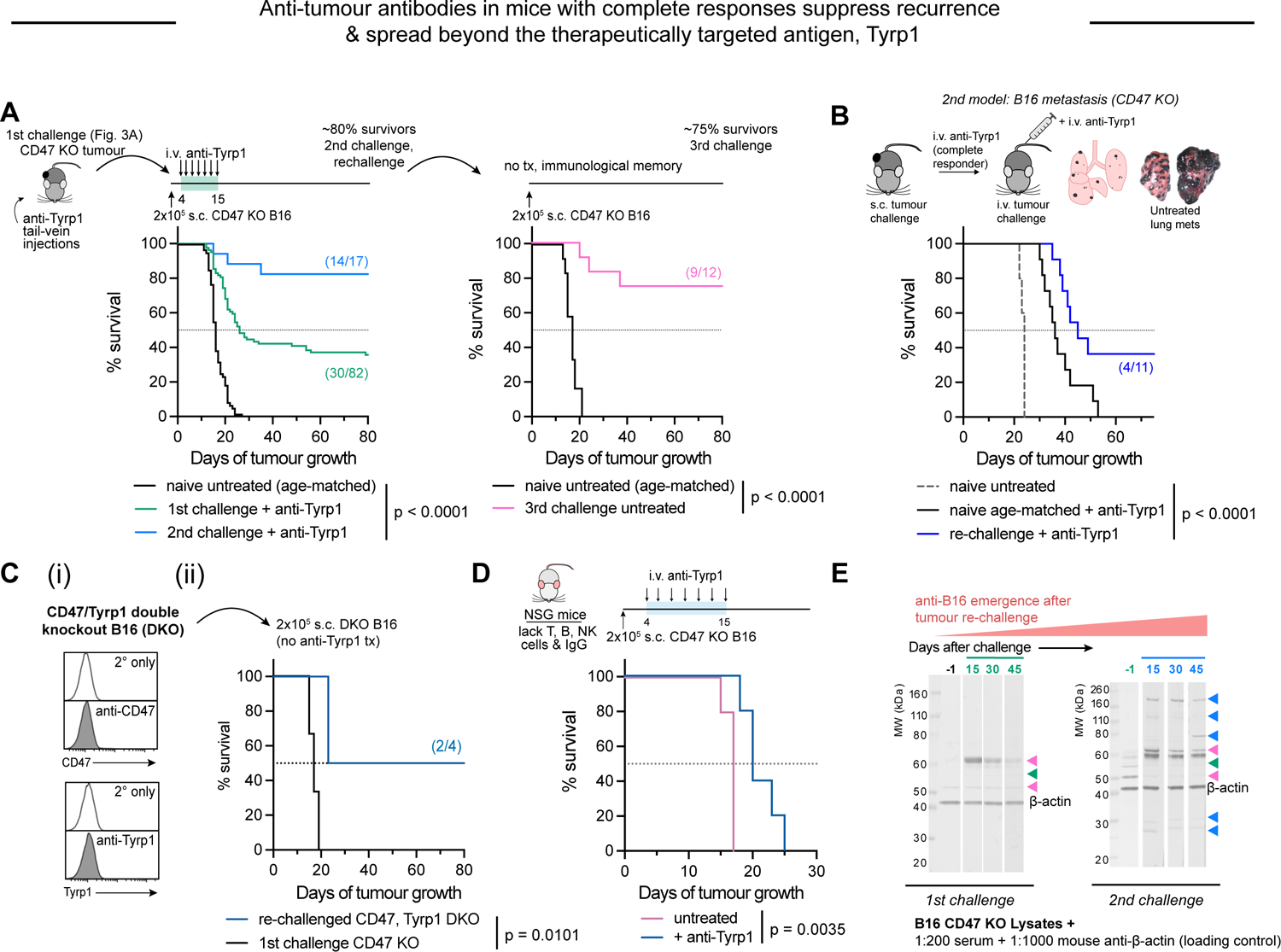
Anti-tumour antibodies in mice with complete responses suppress recurrence & spread beyond the therapeutically targeted antigen, Tyrp1. **A** Treatment scheme of B16 tumours with i.v. tail-vein injections of anti-Tyrp1 (top). Survival curves (bottom) of mice challenged s.c. with 2×10^5^ CD47 KO B16 cells and treated with 250 µg anti-Tyrp1 i.v. (n = 82, from Figure 2C-ii) on days 4, 5, 7, 9, 11, 13, and 15 after engraftment or left untreated (n = 60). Surviving mice challenged with CD47 KO a 2^nd^ time and treated with the same anti-Tyrp1 regimen survive at a rate 2.3-fold higher than age-matched naïve mice (n = 17 across 4 independent experiments) while 75% of mice challenged a 3^rd^ time survive without further treatment (n = 12 per group across 3 independent experiments). Statistical significance was determined by the Log-rank (Mantel-Cox) test. **B** Survival curves of mice bearing experimentally-induced CD47 KO B16 lung metastases. Anti-Tyrp1 delays survival but fails to eliminate these metastases in mice (n = 11 across 2 independent experiments) unless mice had previously eliminated subcutaneous KO tumours (n = 11 across 2 independent experiments). Untreated lung metastasis-bearing mice, n = 5. Statistical significance was determined by the Log-rank (Mantel-Cox) test. **C (i)** Flow histograms of Tyrp1, CD47 KO cells (double knockout, DKO) showing complete depletion of the monoclonal antibody therapeutically-targeted B16 antigen. **(ii)** Survival curves of DKO tumour-bearing mice that had previously survived CD47 KO B16 challenge (+ anti-Tyrp1). 50% of mice survive (n = 4) without additional treatment, while naïve mice challenged with CD47 KO tumours fail to survive (n = 3). Statistical significance was determined by the Log-rank (Mantel-Cox) test. **D** Treatment scheme (top) and survival curves (bottom) of NSG mice engrafted with CD47 KO tumours and treated with anti-Tyrp1 i.v. (n = 5) on days 4, 5, 7, 9 11, 13, and 15 or left untreated (n = 5). Anti-Tyrp1 treatment modestly delayed tumour progression but failed to induce complete anti-tumour responses. Statistical significance was assessed by the Log-rank (Mantel-Cox) test. **E** Anti-B16 IgG emerges after tumour challenge and subsequent re-challenge. Western blotting of CD47 KO B16 lysate with serum as primary probe followed by anti-mouse IgG [H+L] secondary staining shows binding to proteins beyond Tyrp1 that increases in the 2^nd^ tumour challenge.

**Figure 5:**
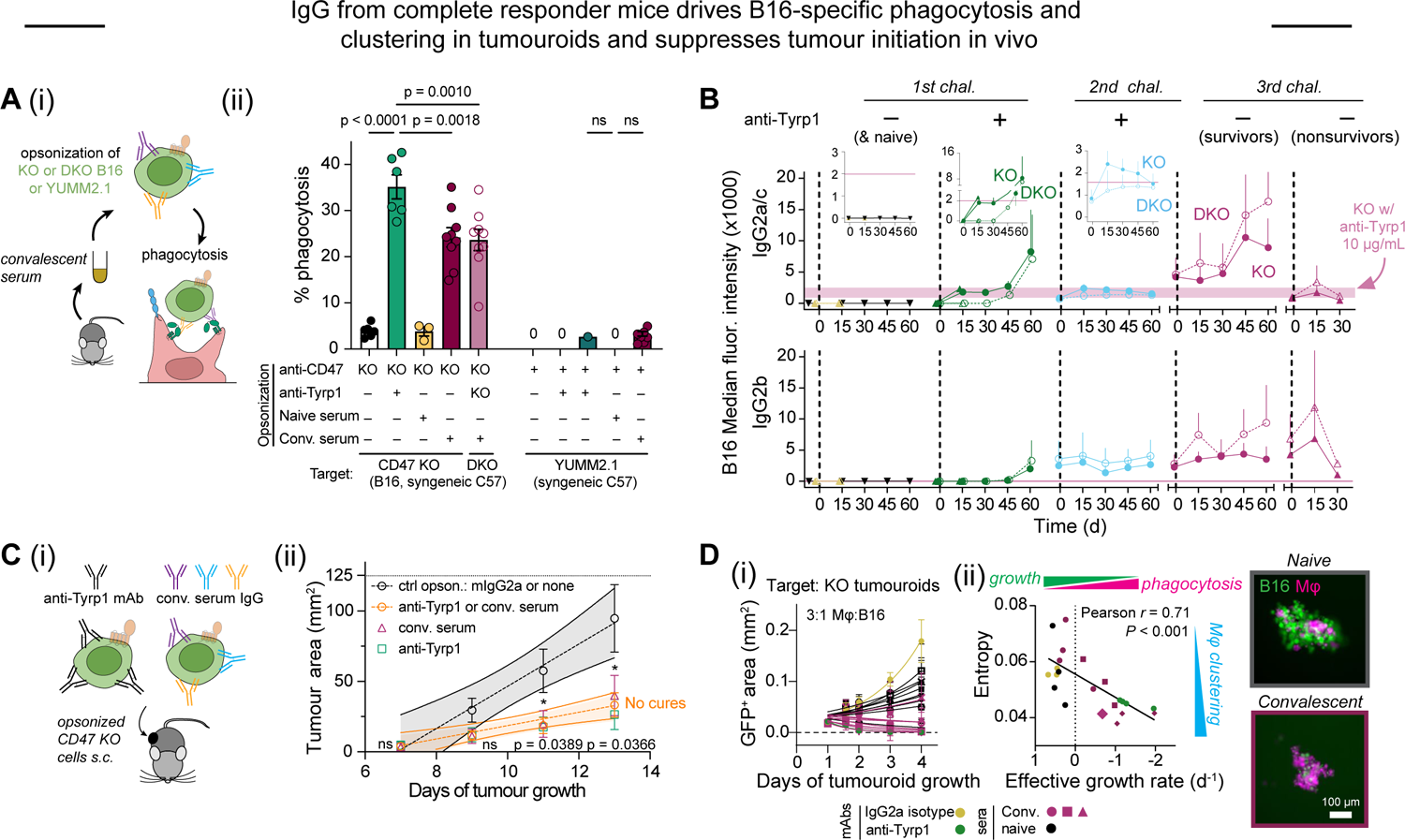
Convalescent serum IgG drives B16-specific phagocytosis and clustering in tumouroids, and suppresses tumour initiation in vivo. **A (i)** Experimental scheme of phagocytosis assay with serum-opsonized DKO B16s or wild-type YUMM2.1s. Convalescent serum collected from mice during and after tumour challenge contains polyclonal anti-B16 that bind and functionally opsonize cells **(ii)** Phagocytosis of serum-opsonized CD47 KO B16s, DKO B16s, or wild-type YUMM2.1s. Convalescent serum IgG from complete responder mice opsonizes B16s and drives phagocytosis even when Tyrp1 is deleted, but does not do so against syngeneic YUMM2.1 cells that share a C57 antigen repertoire. Statistical significance was assessed by one-way ANOVA with Sidak’s test for multiple comparisons, ns not significant (mean ± SEM, opsonization conditions: n = 9 convalescent sera each, n = 3 naïve sera, n = 6 anti-Tyrp1 (B16), n = 3 anti-Tyrp1 (YUMM2.1), n = 6 unopsonized). **B** Kinetic profiles of anti-B16 IgG levels in convalescent sera across the three tumour challenges or in naive sera. Binding of IgG2a/c and IgG2b to CD47 KO (filled symbols) or DKO (open symbols) was measured with subclass-specific secondary antibodies and is reported as the median fluorescence intensity (mean ± SEM, n = 4 survivors, n = 3 non-survivors, n = 5 naive). **C (i)** Schematic of pre-opsonization with serum or anti-Tyrp1 prior to tumour induction. Convalescent sera are incubated with CD47 KO B16 just prior to s.c. flank injection. **(ii)** Tumour growth curves at early timepoints where growth is still in the linear regime. Linear fits show significant growth suppression by CD47 KO B16 cells pre-opsonized with either anti-Tyrp1, convalescent serum, isotype IgG2a, or lacking opsonization (ctrl opsonization n = 9, serum or anti-Tyrp1 opsonization n = 5 per group). Statistical significance at each timepoint between ctrl (black) and combined opsonization (orange) was assessed by two-tailed unpaired t-test with Welch’s correction after significant F test to compare variances, ns not significant. **D (i)** Convalescent sera repress KO tumouroid growth and drive macrophage clustering. Convalescent serum or anti-Tyrp1 can eliminate tumouroids within days whereas naïve serum or isotype control Ab do not. Solid lines are nonlinear regression of the data to a simple exponential of the form *A*(*t*) = *A*_1_ exp[*k_eff_*(*t*-1)] (mean ± SEM, n = 3 or 4 tumouroids per sample for 9 convalescent and 9 naïve sera). **(ii)** Entropy measured by image file compression (Fig. 2A) correlated (Pearson *r* = 0.71, p = 0.0003) with the effective growth rate. For simplicity, data are shown for convalescent sera that produced high (circles), medium (squares), and low (diamonds) entropy clusters (and low, medium, and high growth repression, respectively) and for a representative naïve serum sample (n = 3 or 4 tumouroids per serum or antibody condition). Representative fluorescence images depict high and low entropy macrophages, respectively, in tumouroids on day 2, ∼24 h after addition of macrophages and naïve (top) or convalescent serum (bottom). Scale bar: 100 µm.

### Increasing phagocytic macrophage density eliminates tumours

To assess whether conditions of maximum phagocytosis – including high density of macrophages – are effective *in vivo,* CD47 KO tumours were established in mice (for 4 days) and then given one intravenous (i.v.) injection of anti-Tyrp1 (**Fig 2B-i**). Mice were sacrificed 24 h later, and compared to control, the monoclonal antibody caused 3-fold smaller tumours and 3-fold more macrophages (F4/80+ cells) in the tumour (**Fig 2B-ii-iv**). The remaining immune cells (CD45+, F4/80-) did not differ between conditions. Macrophages also segregated within treated tumours and co-localized with B16-produced melanin (**Fig 2B-ii,v**), which is consistent with observations of tumouroids (Extended Data Fig. 2F).

Longer duration experiments showed anti-Tyrp1 i.v. injections eliminated CD47 KO tumours in ∼40% of mice, whereas various control tumours, including untreated CD47 KO and both treated or untreated WT tumours all showed the expected exponential growth (**Fig 3A-i,ii**). The WT results are consistent with recent studies^26^, and control IgG2a antibody or vehicle injections also showed no effect (Extended Data Fig. 6). To begin to assess the effect of increased macrophage numbers, fresh bone marrow cells (containing 5-10% monocytes and macrophages) were also i.v. injected together with anti-Tyrp1. The combination eliminated CD47 KO tumours in ∼80% of mice (**Fig 3A-ii**).

**Figure 6:**
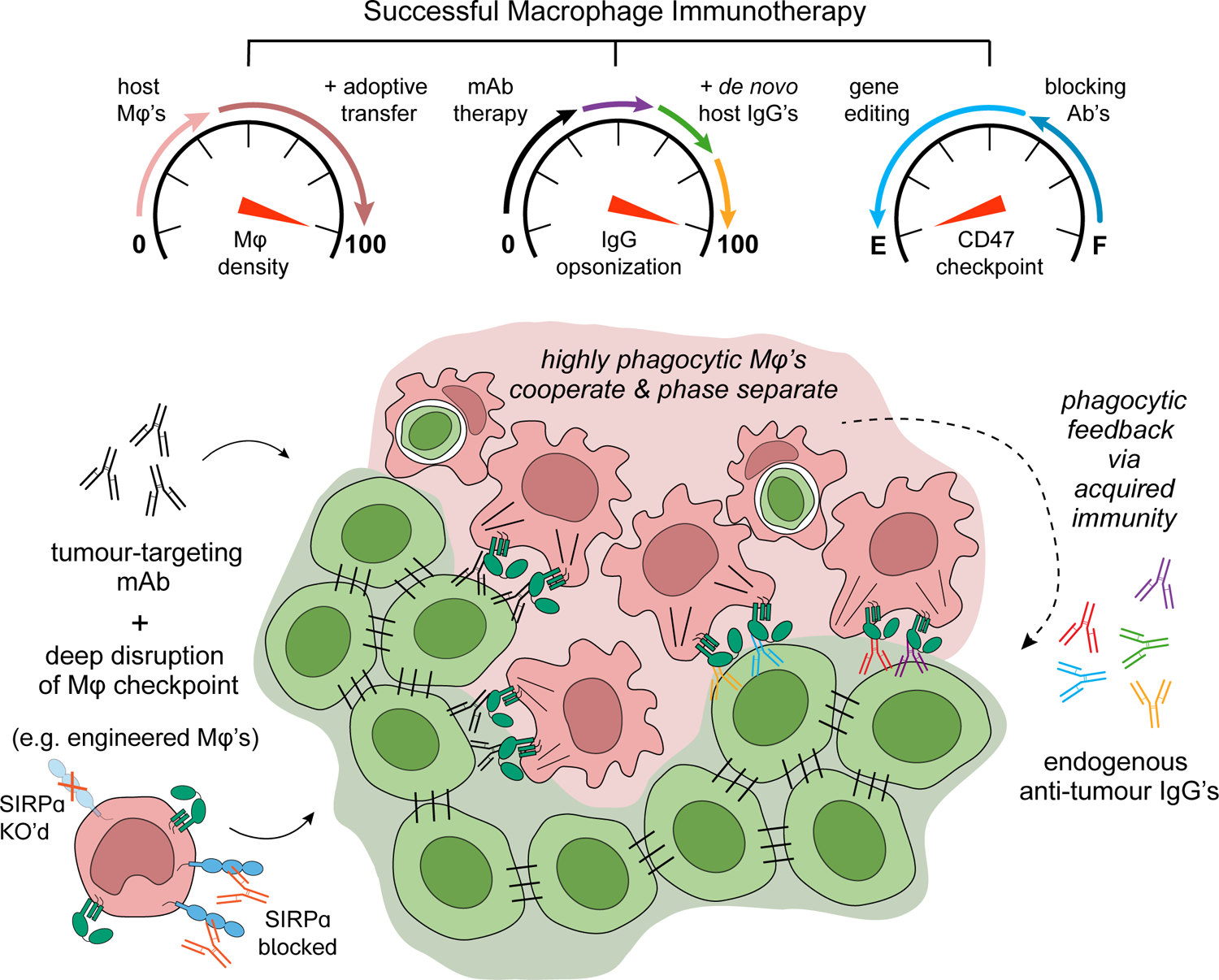
Phagocytic feedback in tumour elimination by cooperative macrophages. Macrophage density and cooperativity, IgG-opsonization, and efficient blockade of the macrophage checkpoint underlie successful macrophage immunotherapy of solid tumours. Phase separating macrophages cooperatively engulf cancer cells when CD47 signaling is disrupted deeply and cancer cells are opsonized with IgG to maximize engulfment. Macrophage phagocytosis in tumours treated with anti-tumour IgG monoclonal antibody (mAb) ultimately leads to the production of *de novo* endogenous anti-tumour IgG that provide phagocytic feedback.

Given that WT tumours are unaffected by i.v. injections of anti-CD47 and anti-Tyrp1, we hypothesized that we could eliminate WT tumours by adding i.v. injections of marrow engineered by addition of anti-SIRPα to block CD47 interactions (A’PB cells in **Fig 3B**). This strategy enabled high levels of phagocytosis and eliminated tumouroids *in vitro* at high macrophage:B16 ratios (**Fig 3C-i**, Extended Data Fig. 7A). Even *in vivo* without systemic anti-CD47, we found complete responses in ∼10% of mice with WT tumours (Extended Data Fig. 7B) – which increased to ∼17% with anti-CD47 injections and to ∼38% with a second i.v. injection of the marrow (**Fig 3C-ii**) (Extended Data Fig. 7B). Such marrow macrophages isolated from tumours have been found to exhibit markers consistent with phagocytic activity^11^. To confirm macrophages are the effector cells and to potentially improve upon antibody-based blockade with a genetic engineering approach, we deleted the SIRPα gene in conditionally immortalized macrophage (CIM) progenitors that retain macrophage differentiation ability and phenotypes^35, 36^ (Extended Data Fig. 8). These engineered macrophages also phagocytosed target B16s highly with anti-Tyrp1 opsonization *in vitro* (**Fig 3D-i**). Injection of these cells i.v. with anti-Tyrp1 (but no added anti-CD47) produced complete anti-tumour responses against WT tumours in ∼60% of mice (**Fig 3D-ii**). Thus, as we observed with the immuno-tumouroid results, a succinct summary of many experiments shows that maximizing phagocytosis is necessary for tumour elimination and high numbers of phagocytic macrophage maximize efficacy (**Fig 3E**, Extended Data Fig. 7C).

Macrophages and related phagocytic immune cells provide just a first line of defense but often initiate acquired immunity^24, 25^. We therefore challenged surviving mice with a second injection of CD47 KO cells 80 days after the initial challenge and again treated with anti-Tyrp1 (a prime-boost strategy for an anti-cancer vaccine). From the initial cohort in which the complete response rate was about 40% (**Fig 3A-ii**), about 80% of the re-challenged mice survived (**Fig 4A**). Age-matched naïve mice receiving their first challenge with or without treatment responded similarly to the younger cohorts. A third tumour challenge was left untreated (i.e. no anti-Tyrp1), and 75% again resisted tumour growth, which indicates a durable immunological memory. Whitening of the melanized fur was evident across all three challenges of treated mice in the KO cohorts, consistent with an immune response against normal melanocytes that occurs over time with anti-Tyrp1 as mediated by complement or Fc-receptors^37^. A second model of B16s involving metastasis to lung with CD47 KO cells showed anti-Tyrp1 prolongs survival of naïve mice until all mice succumb with clinical symptoms (**Fig 4B**). Importantly, however, mice that previously showed complete responses against subcutaneous tumours *could* achieve a complete anti-tumour response against lung metastasis when treated with anti-Tyrp1, as demonstrated by long-term survival of ∼35% of mice.

### Convalescent serum IgG drives tumour phagocytosis & macrophage clustering

Despite the importance of anti-Tyrp1 for CD47 KO tumour elimination, this monoclonal antibody was *not* used in a third challenge of CD47 KO tumours cells in which 75% of mice resisted the tumour (**Fig 4A**). We hypothesized therefore that an acquired immune response is generated that might also extend beyond Tyrp1. We thus knocked out Tyrp1 in the CD47 KO cells (double knockout, DKO) and engrafted these DKO cells in complete responders from the initial CD47 KO treatment cohort. We found that without monoclonal anti-Tyrp1 treatment, which would not be effective against DKO tumours anyway, 50% of mice survived (**Fig 4C-i,ii**). Although this result provided initial evidence of acquired immunity unrelated to Tyrp1, we tested the need for an acquired immune system by assessing tumour growth in NSG mice that lack all adaptive immunity (no antibodies or T, B, or NK cells). Growth of CD47 KO B16 tumours proves similar in NSG mice to growth in C57 mice. Although i.v. injections of anti-Tyrp1 slow the growth of tumours in NSG mice, no NSG mice survived beyond 25 days (**Fig 4D**) in contrast to the 40% long-term survivors among immunocompetent C57 mice under the same conditions (**Fig 3A-ii**). Macrophages in NSG mice display Fc-receptors for binding IgG^11^, and we have shown CD47-knockdown human tumours in NSG mice can be suppressed with i.v. injections of anti-human monoclonal antibody. However, the lack of complete anti-tumour responses in the previous study and again here with B16 tumours implicates a role for acquired immunity.

Serum collected throughout tumour challenge and treatment experiments was used to immunoblot B16 lysates, which revealed an increasing number of bands with progression of challenges in support of our hypothesis of immune responses against B16 antigens beyond Tyrp1 (**Fig 4E**). To test whether such serum antibodies are functional to promote phagocytosis, we added serum from various mice to CD47 KO and DKO B16 cells and then added this mixture to macrophages in culture. Most convalescent sera increased phagocytosis relative to naïve serum (**Fig. 5Ai-ii**, Extended Data Fig. 9B), with similar results for CD47 KO cells, WT B16 parental cells, and sub-clones as targets for phagocytosis as well as for surface binding (Extended Data Fig. 9A,C,D). As expected, anti-Tyrp1 does not drive phagocytosis of DKO cells, and neither anti-Tyrp1 nor the serum from complete responders has any effect on engulfment of YUMM2.1 melanoma cells that were also derived from C57 mice.^38^ YUMM2.1 cells generate tumours in mice that previously eliminated B16 tumours, even when treated with SIRPα-blocked marrow macrophages (Extended Data Fig. 9E). Convalescent serum antibodies thus target B16-specific antigens beyond Tyrp1 but not antigens expressed by a syngeneic melanoma or xenogeneic antigens present on cultured cell lines (e.g., bovine antigens from media containing fetal bovine serum). Assays of IgG subclasses confirm the presence of pro-phagocytic IgG2a/c and IgG2b^39^ that bind CD47 KO cells and DKO cells after second and third challenges (**Fig 5B**), with other IgG subtypes also detected (Extended Data Fig. 10A-C). Further consistent with a progressive prime-boost vaccination and antigenic spread beyond Tyrp1, survivors of the *third* B16 challenge (with no monoclonal anti-Tyrp1 injected) showed higher levels of IgG2a/c compared to non-survivors. To test the function of this serum *in vivo*, CD47 KO B16s were opsonized with the serum just prior to subcutaneous implantation in naïve mice (**Fig 5C**). Tumour growth was suppressed by days 11 and 13 similar to anti-Tyrp1 and relative to controls. However, because any pre-bound IgG will be diluted by B16 proliferation or else lost by dissociation, we assessed complete elimination of tumour cells by convalescent serum and macrophages with tumouroids.

Convalescent serum IgG added with macrophages to CD47 KOs tumouroids eliminated tumouroids for the most potent samples (**Fig 5D-i**). Importantly, serum activity against tumouroids correlated with macrophage clustering at day 2 similar to anti-Tyrp1 (**Fig 5D-ii**), consistent with effects of monoclonal anti-Tyrp1 (Fig 2A). Serum alone had no effect (Extended Data Fig. S10D), indicating macrophages are the effector cells for the anti-B16 serum IgG.

## Discussion

Macrophage immunotherapy of solid tumours benefits from (1) macrophage number and cooperativity, (2) tumour opsonization with anti-cancer IgG that activates Fc receptors, and (3) inhibition of the macrophage checkpoint SIRPα that binds CD47 (**Fig 6**). Maximizing *all three factors* can eliminate established syngeneic B16 tumours in immunocompetent mice and initiate *de novo* anti-tumour IgG in a vaccine-like response to monoclonal antibody therapy. There is already precedent for anti-RBC IgG being induced with CD47 depletion in mouse^40^, but this phenomenon is otherwise understudied. Isolation and cloning of the new tumour-specific pro-phagocytic IgG induced after CD47-disruption can help address a critically important need for functional monoclonal IgG’s that can be combined with clinically relevant CD47 blockade for anti-tumour efficacy. Thus, while the biopharmaceutical industry is expert at the cloning and manufacturing of antibodies, our approach is tailored to initial identification of tumour-opsonizing IgG that synergize with macrophage checkpoint disruption.

The most promising clinical application of CD47-SIRPα blockade to date combines anti-CD47 (magrolimab) and anti-CD20 (rituximab) against a liquid tumour, non-Hodgkin’s lymphoma^15^, but B cell depletion by this clinical treatment limits the capacity of patients to develop any anti-tumour IgG as described here, which thus prevents a phagocytic feedback. Anti-cancer IgG have been identified in cancer patients, including anti-(human Tyrp1) from a melanoma patient^41, 42^, but a monoclonal antibody that targets Tyrp1 is not sufficient on its own for clinical efficacy against melanoma^32^. Our analysis of the human metastatic melanoma cohort in TCGA reveals that high *TYRP1* expression is associated with worse survival than low *TYRP1*, independent of prior treatment of patients (Extended Data Fig. S11) and indicates that TYRP1 remains targetable even in aggressive human melanomas. Low *CD47* expression in the same patients also shows worse survival than high *CD47*, despite opposite trends for some other cancers^12, 43^. Low CD47 can nonetheless be protective as demonstrated by the failure of macrophages to eliminate tumouroids at anti-CD47 concentrations < 100 nM (Fig 1F-ii).

The important but under-appreciated role for the number and cooperativity of *phagocytic* macrophages in solid tumours does not contradict the frequent observation that macrophages are abundant and provide a poor prognosis in human primary and metastatic tumours including melanoma^44^. Tumour-associated macrophages are generally not phagocytic^45^, although phagocytic macrophage aggregates have been reported in thyroid cancer and associate with decreased risk of metastasis^46^. Moreover, the negative correlation between macrophage abundance and survival in follicular lymphoma patients receiving chemotherapy was abrogated and potentially even reversed in patients receiving rituximab and chemotherapy combination^47, 48^. Therefore, the role of macrophages in tumour progression seems dependent on the physical state of the macrophages and patient treatment status.

Considerations of the physical properties of the tumour microenvironment including its immune cell residents^4^ are critical for success of immunotherapies that rely for efficacy on (1) IgG permeation, including CD47-blocking monoclonal antibodies and (2) immune cell infiltration and segregation into phagocytic clusters. With respect to blocking antibodies, we did find that anti-SIRPα on macrophages immobilizes SIRPα rather than directly blocking CD47 interactions (Extended Data Fig. 12), which indicates a biophysical mechanism beyond mere antagonism of a molecular interaction. Our SIRPα-blocked monocytes/macrophages take advantage of the ability of such cells to actively migrate and permeate through very tight spaces^11^, overcoming limits of passive permeation. Complete knockout of SIRPα in CIM’s improved responses beyond those achieved with antibody blockade on adoptively transferred marrow cells. Further engineering of macrophages and new strategies might also prove beneficial^49, 50^. Regardless, durability against loss of epitopes, resistance, and metastasis might thus result from the emergent phagocytic feedback.

## Materials and Methods

### Cell culture

B16-F10 (CRL-6475) cells were obtained from American Type Culture Collection and cultured at 37 °C, 5% CO_2_ in either RPMI-1640 (Gibco 11835-030) or Dulbecco’s Modified Eagle Medium, (DMEM, Corning 10-013-CV) supplemented with 10% (v/v) fetal bovine serum (FBS, Sigma F2442), 100 U/mL penicillin and 100 µg/mL streptomycin (1% P/S, Gibco 15140122). B16 KO cell lines were generated as described previously^1^ using single guide RNA (sgRNA) constructs targeting CD47 (5’-TCCCCGTAGAGATTACAATG) and Tyrp1 (5’-CTTGTGGCAATGACAAATTG) and cultured under the same conditions as the parental wild-type B16-F10 cell line. YUMM2.1 cells were a gift from Dr. Chi Van Dang at the Wistar Institute and were cultured under the same conditions in DMEM.

### Antibodies

Antibodies used for *in vivo* treatment and blocking, *in vitro* phagocytosis, flow cytometry, and Western blotting are as follows: anti-mouse/human Tyrp1 clone TA99 (BioXCell BE0151), isotype IgG2a control clone C1.18.4 (BioXCell, BE0085), anti-mouse CD47 clone MIAP301 (BioXCell BE0270), isotype control rat IgG2a (BioXCell, BE0089), anti-mouse SIRPα clone P84 (BioLegend 144004), isotype control rat IgG1 (BioLegend, 400414). Low-endotoxin and preservative-free antibody preparations were used for *in vivo* opsonization and blocking and *in vitro* phagocytosis experiments. The following BioLegend antibodies were used for flow cytometry staining: Pacific Blue anti-Ly6C clone HK1.4 (128013), PE anti-Ly6C clone HK1.4 (128007), PE anti-F4/80 clone BM8 (123109), APC/Cy7 anti-F4/80 clone BM8 (123118), Brilliant Violet 605 anti-CD11c (117334), PerCP/Cy5.5 anti-mouse CD45.2 clone 104 (109827), APC anti-mouse CD45.2 clone 104 (109814), PE/Cy7 anti-mouse/human CD11b clone M1/70 (101216), FITC anti-mouse SIRPα clone P84 (144006), APC/Cy7 anti-mouse IgG1 clone RMG1-1 (406619), APC anti-mouse IgG2a clone RBG2a-62 (407110), PE/Cy7 anti-mouse IgG2a clone RBG2a-62 (407113), APC anti-mouse IgG2b clone RMG2b-1 (406711), PE/Cy7 anti-mouse IgG2b clone RMG2b-1 (406714), biotin anti-mouse IgG3 clone RMG3-1 (406803), PE streptavidin (405203), Alexa Fluor (AF) 647 donkey anti-mouse IgG (ThermoFisher A31571), AF647 goat anti-rat IgG (ThermoFisher A21247). Primary antibodies used in Western blotting or immunofluorescence (IF) microscopy were anti-β-actin clone C4 (Santa Cruz sc 47778), mouse anti-lamin A/C clone 4C11 (Cell Signaling Technology 4777) rabbit anti-lamin B1 (Abcam ab16048), mouse anti-N-cadhein clone 13A9 (Biolegend 844701), and mouse anti-α-tubulin (Sigma T9026). Secondary antibodies used in Western blotting and IF were HRP sheep anti-mouse IgG (GE Life Sciences NA931V), goat anti-mouse IgG IRDye800CW (LiCoR 926-32210), AF488 donkey anti-mouse IgG (ThermoFisher A21202), and AF647 donkey anti-rabbit IgG (ThermoFisher A31573).

### Mice

C57BL/6J mice (Jackson Laboratory 000664) were 6-12 weeks old at the time of first challenge unless otherwise specified. Age-matched mice were used as controls in rechallenge experiments. NOD-*scid* IL2Rγ^nul^ (NSG) mice aged 6-12 weeks old were obtained from the Stem Cell & Xenograft Core at the University of Pennsylvania. All experiments were performed in accordance with protocols approved by the Institutional Animal Care and Use Committee of the University of Pennsylvania.

### Bone marrow-derived macrophages (BMDMs)

Bone marrow was harvested from donor mice, lysed with ACK buffer (Gibco A1049201) to deplete red blood cells, and cultured on Petri dishes for 7 days in Iscove’s Modified Dulbecco’s Medium (IMDM, Gibco 12440053) supplemented with 10% FBS, 1% P/S, and 20 ng/mL recombinant mouse macrophage colony-stimulating factor (M-CSF, BioLegend, 576406). Successful differentiation was confirmed by flow cytometry staining with antibodies against macrophage markers. Cytokine-primed BMDMs were cultured in RPMI growth media + 20 ng/mL M-CSF + 20 ng/mL IFNγ (BioLegend 575302) or 20 ng/mL IL-4 (BioLegend 574302) for 48 h prior to use in tumouroid assays or prior to analysis of protein expression by immunofluorescence microscopy or flow cytometry. For immunofluorescence microscopy, differentiated BMDMs were detached and re-plated on bare glass coverslips at a density of ∼5.5 x 10^3^ per cm^2^ in RPMI growth media + 20 ng/mL M-CSF. Cytokines were added 3 h later when cells were mostly attached to the glass.

### Conditionally immortalized macrophage (CIM) progenitors

Macrophage progenitors were generated as previously described^2,3^. CIMs were cultured in suspension in 10 cm Petri dishes in RPMI-1640 (A1049101, ThermoFisher) supplemented with 10% FBS and 1% P/S as above. Media was also supplemented with 43 µM β-estrogen (Sigma Aldrich, E2758-250MG) and 10 ng/mL recombinant mouse GM-CSF (BioLegend, 576302). Cells were passaged every 2 days at sub-confluent concentration of 1×10^5^ cells/mL. To differentiate for phagocytosis assays, cells were washed twice with 5% FBS/PBS to remove excess β-estrogen and plated in supplemented DMEM (10% FBS/1%PS) with 20 ng/mL recombinant mouse M-CSF for 7 days.

### *In vitro* phagocytosis

For 2D phagocytosis assays, BMDMs were detached and re-plated in 24 well plates at a density of 1.8 ×10^4^ per cm^2^ in IMDM supplemented with 10% FBS, 1% P/S, and 20 ng/mL M-CSF. The next day, BMDMs were labeled with 0.5 µM CellTracker Deep Red dye (Invitrogen C34565) according to the manufacturer’s protocol. Following staining, macrophages were washed and incubated in serum-free IMDM (i.e., IMDM supplemented with 0.1% (w/v) bovine serum albumin and 1% P/S). In some experiments, target B16 cell lines were labeled with carboxyfluorescein diacetate succinimidyl ester (Invitrogen V12883) also according the manufacturer’s protocol. B16 were detached and opsonized with 10-20 µg/mL anti-Tyrp1, with mouse IgG2a isotype control antibody, or with 5% (v/v) mouse serum collected during the course of tumour challenge as described below in serum-free IMDM for 30 min on ice. For CD47 blockade experiments, 20 µg/mL MIAP301 or rat IgG2a isotype control antibody was added during opsonization. Opsonized B16 suspensions were added to BMDMs at a ∼2:1 ratio and incubated at 37 °C, 5% CO_2_ for 2 h. Nonadherent cells were removed by washing with PBS and remaining cells were fixed with 4% formaldehyde, counterstained with Hoechst 33342 (Invitrogen H3570), and imaged on an Olympus IX inverted microscope with a 40x/0.6 NA or 20x/0.4 NA objective.

The Olympus IX microscope was equipped with a Prime sCMOS camera (Photometrics) and a pE-300 LED illuminator (CoolLED) and was controlled with MicroManager software v1.4 or v2^4^.

### Tumouroids

To generate surfaces conducive to B16 tumouroid formation, 96-well plates (Greiner Bio-One 650161) were either coated with 70 µL of 2% agarose in water or PBS or treated for 20 min with 100 µL of anti-adherence rinsing solution (StemCell Technologies 07010). The wells were washed with PBS, and then blocked with RPMI containing 0.5-1% bovine serum albumin (BSA, Sigma) 37 °C for 1 h. B16 were detached by brief trypsinization, resuspended in RPMI growth media (RPMI supplemented with 10% FBS, 1x non-essential amino acid solution [Gibco 11140050], 1 mM sodium pyruvate [Gibco 11360070], and 50 µM β-mercaptoethanol [Gibco 21985023]) at a concentration of 1×10^3^ or 1×10^4^ per mL, and added to passivated wells in a volume of 100 µL. A single aggregate of B16 cells formed in each well within 12 h. For phagocytosis versus growth studies, CellTracker Deep Red-labeled BMDMs and antibodies (final concentration 20 µg/mL anti-Tyrp1 and/or 20 µg/mL anti-CD47 or 10 µg/mL anti-SIRPα) or mouse serum (final concentration 1:200 convalescent or naïve) were added 24 h later in a volume of 20 µL RPMI growth media with 120 ng/mL M-CSF. For myosin II inhibition studies, 20 µM blebbistatin (MilliporeSigma 203389) or an equal volume of DMSO vehicle was added to B16 tumouroids for 1 h prior to addition of BMDMs. Tumouroids were imaged on the Olympus IX microscope with 10x/0.3 NA or 4x/0.13 NA objectives. Tumouroid areas were measured in ImageJ after thresholding GFP fluorescence using MaxEntropy or Li algorithms and manually adjusting the threshold as needed.

For confocal imaging, clusters were fixed ∼20 h after addition of BMDMs and antibodies by adding an equal volume of 4% formaldehyde directly to the media in each well for 30 min at room temperature. Half of this solution was removed and replaced with an equal volume of 2% formaldehyde for 30 min. The clusters were washed with PBS to remove fixative repeating the procedure of removing and replacing half the volume at each step. The clusters were transferred via a wide-bore pipette tip to an 8-well chambered coverglass (Lab-Tek) and counterstained with 5 µg/mL Hoechst 33342 (Thermo). Confocal images were acquired on a Leica TCS SP8 with a 20x/0.75 NA objective. Radial fluorescence profiles were calculated in ImageJ by thresholding max intensity projections of GFP and Deep Red fluorescence and calculating the tumouroid centroid, which was used as the input for *r* = 0 in the Radial Profile ImageJ plugin applied individually to the GFP and Deep Red channels. This plugin calculates the average fluorescence intensity *I*(*r*) for a circle of radius *r* centered at the inputted position of the tumouroid centroid. *I*(*r*) values were normalized by dividing by the total average fluorescence intensity Σ*I*(*r*) for all *r*.

### Entropy image analysis

Fluorescence images of CellTracker Deep Red-labeled macrophages were converted to binary images of black cells on a white background using the Otsu threshold algorithm in ImageJ. The number of black pixels was determined from the image histogram, and the image was saved as a PNG file, which is a form of lossless file compression to determine the compressed file size as a measure of entropy. Random images were generated and analyzed using the Python NumPy and Pillow packages, respectively.

### Transcriptomic analysis

Public microarray data for BMDMs treated with IFNγ for 18 h versus control BMDMs (GSE60290^5^) and for BMDMs treated with IL-4 for 24 h versus control BMDMs (GSE69607^6^) were accessed and analyzed using the GEO2R interactive web tool (https://www.ncbi.nlm.nih.gov/geo/geo2r/). Differential expression was considered significant for p < 0.05, where the adjusted p-values were computed by the default method in GEO2R that uses the false discovery rate method of Benjamini and Hochberg.

### Flow cytometry

Phagocytosis of B16 by macrophages in tumouroids was assessed by pooling identically treated tumouroids from a single 96-well plate, disaggregating them to single cell suspensions, and analyzing cell suspensions by flow cytometry. Tumouroids were collected into FACS buffer (PBS plus 1% BSA and 0.1% sodium azide) ∼18 h after addition of BMDMs ± anti-Tyrp1, pelleted by centrifugation and resuspended in FACS buffer + 0.5 mM EDTA for 15-30 min at room temperature. The suspension was pipetted up and down until the cells were deemed to be disaggregated by inspection on a hematocytometer. Viable cells were distinguished by staining with Zombie aqua fixable viability dye (BioLegend 423101) in PBS for 15 min at room temperature. The cells were washed with FACS buffer, fixed with 4% formaldehyde for 15 min at room temperature, and stored at 4 °C until analysis. Flow cytometry was performed on a BD LSRII (Benton Dickinson) and data were analyzed with FCS Express 7 software (De Novo Software). Doublets, cell debris, and nonviable cells were excluded and GFP^+^ DeepRed^+^ events were considered to be phagocytic macrophages.

Bone marrow-derived macrophages (BMDMs) were differentiated and primed as described above. Fc-receptors were blocked with rat anti-mouse CD16/32 (clone 2.4G2, BD 553149) and rat anti-mouse CD16.2 (clone 9E9, BioLegend 149502) prior to staining with macrophage marker antibodies.

For serum binding analyses, B16 or YUMM2.1 cells were detached by brief trypsinization, washed, and resuspended in FACS buffer containing primary antibody or 5% (v/v) mouse serum collected as described below. Cell suspensions were incubated at 4°C for 30 min and agitated to prevent cell settling. Cells were washed three times with FACS buffer and incubated with fluorophore-conjugated secondary antibodies in FACS buffer for 30 min at 4°C and later with PE-streptavidin if required to label biotinylated anti-mouse IgG3. Finally, cells were washed three times and resuspended in FACS buffer containing 0.2 µg/mL DAPI (Cell Signaling Technology). For samples that could not be analyzed on the day of staining and required fixation, cells were stained with Zombie aqua fixable viability dye in PBS for 15 min at room temperature prior to staining with antibodies or serum, and DAPI was omitted from the final fixed cell suspension. Those samples were fixed with FluoroFix Buffer (Biolegend 422101) or 2-4% paraformaldehyde. For experiments analyzed in the same plot, the same lots of primary and secondary antibodies and identical staining conditions were used. When these analyses were performed on different days, UltraRainbow calibration beads (Spherotech URCP-38-2K) were used to adjust the photomultiplier tube voltage in each channel to maintain the median fluorescence intensity of the brightest peak within a tolerance of ± 5%.

### Tumour models

B16 and YUMM2.1 cell lines cultured in DMEM growth media were detached by brief trypsinization, washed with PBS, and resuspended at 2×10^6^ per mL in PBS. Cell suspensions remained on ice until injection. Fur on the injection site (usually the right flank) was wet slightly with a drop of 70% ethanol and brushed aside to visualize the skin. A 100 µL bolus (containing 2×10^5^ tumour cells) was injected beneath the skin. Treated mice received i.v. injections of anti-Tyrp1 clone TA99 (250 µg antibody in 100 µl PBS) via the lateral tail vein on days 4, 5, 7, 11, 13, and 15 post tumour cell inoculation. Where indicated, mice also received i.v. injections of anti-CD47 clone MIAP301 (83 µg antibody in 100 µl PBS dosed in the same volume as anti-Tyrp1). Tumours were monitored by palpation and measured with digital calipers. The projected area was roughly elliptical and calculated as *A* = π/4 x *L* x *W* where *L* is the length along the longest axis and *W* is the width measured along the perpendicular axis. A projected area of 125 mm^2^ was considered to be terminal tumour burden for survival analyses.

### Serum collection

Blood was drawn retro-orbitally and allowed to clot for 30-60 min at room temperature in a microcentrifuge tube. The serum was separated from the clot by centrifugation at 1500 x *g* and stored at −20 °C for use in flow cytometry, phagocytosis assays, and Western blotting.

### Immune infiltrate analysis of tumours

For flow cytometry of immune cell markers in tumours, mice were treated with a single dose of i.v. anti-Tyrp1 at day 4 (96 h) post inoculation and sacrificed 24 h later. Tumours were photographed, then excised and placed into 5% FBS/PBS. Tumours were then disaggregated with Dispase (Corning 354235) supplemented with 3 mg/mL Collagenase Type IV (Gibco 17104019) and DNAseI (Sigma-Aldrich 10104159001) for 30 min at 37 °C, centrifuged for 5 min at 300 x *g*, and resuspended in 1 mL of ACK lysing buffer for 12 min at RT. Samples were centrifuged for 5 min at 300 x *g*, washed with FACS buffer, and incubated in FACS 100 µl FACS buffer containing fluorophore-conjugated antibodies to immune markers on ice for 30 min. Samples were then washed with FACS buffer and fixed with FluoroFix Buffer for 30 min at RT prior to analysis on a flow cytometer.

For IF staining of tumour sections, mice were treated in the same manner. Whole tumours were then excised, fixed in 4% paraformaldehyde overnight at 4 °C, and stored in 70% ethanol. The Comparative Pathology Core (University of Pennsylvania) embedded the tissues in paraffin, sectioned, and stained the tumour sections with anti-F4/80 according to their standardized protocols. Sections were imaged on an Olympus microscope as described above. Radial profile analysis was conducted with the Radial Profile Plot plugin for ImageJ. Sectioning and trichrome staining of B16 tumours were performed by the Molecular Pathology and Imaging Core (University of Pennsylvania). Tile scan images of the trichrome-stained tumour section were acquired on the EVOS FL Auto Imager with a 10x/0.25 NA objective (Thermo Fisher).

### Adoptive cell transfers

Fresh bone marrow was harvested as above through the RBC lysis step. Marrow cells were then counted on a hemocytometer and resuspended at 8×10^7^ cells/mL in 5% FBS/PBS. To block SIRPα, cells were then incubated with anti-SIRPα clone P84 (18 µg/mL) for 45 mins at room temperature on a rotator, centrifuged to remove unbound P84, and re-suspended again at 8×10^7^ cells/mL in 2% FBS/PBS with or without 2.5 mg/mL anti-Tyrp1 (TA99). Marrow cells (2×10^7^ cells in 250 µl 2% FBS/PBS) were injected i.v. into tumour-bearing mice 4 days after tumour engraftment. SIRPα KO CIM progenitors (4×10^6^ cells in 250 µl 2% FBS/PBS with 2.5 µg/mL anti-Tyrp1) were injected i.v. in the same manner. YUMM2.1 tumour-bearing mice received a single dose of 2×10^7^ SIRPα-blocked marrow cells i.v. on day 4.

### Western blotting

Lysate was prepared from B16 CD47 KO cells using RIPA buffer containing 1x protease inhibitor cocktail (Sigma P8340) and boiled in 1x NuPage LDS sample buffer (Invitrogen NP0007) with 2.5% v/v β-mercaptoethanol. For detection of N-cadherin, B16 proteins were fractionated by ultracentrifugation. Proteins were separated by electrophoresis in NuPAGE 4-12% Bis-Tris gels run with 1x MOPS buffer (Invitrogen NP0323) and transferred to an iBlot nitrocellulose membrane (Invitrogen IB301002). The membranes were blocked with 5% nonfat milk in Tris buffered saline (TBS) plus Tween-20 (TBST) for 1 h and stained with primary antibodies or with 5% (v/v) mouse serum overnight at 4 °C with agitation. The membranes were washed with TBST and incubated with 1:500 secondary antibody conjugated with horseradish peroxidase (HRP) or with 1:5000 secondary antibody conjugated with IRDye800CW in 5% milk in TBST for 1 h at room temperature with agitation. The membranes were washed again three times with TBST, then TBS. Membranes probed with HRP-conjugated secondary were developed with a 3,3’,5,5’-teramethylbenzidine (TMB) substrate (Genscript L0022V or Sigma T0565). Developed membranes were scanned and analyzed with ImageJ (National Institutes of Health). Membranes probed with IRDye800CW-conjugated secondary were imaged on an Odyssey near-infrared scanner (LiCor).

### Pipette aspiration rheology

Tumouroids were formed in non-adhesive well-plates and transferred via wide-bore pipette to a glass-bottom dish (Mattek). Fresh tumours were harvested and stored in RPMI + 10% FBS. The tumour was held in a chamber consisting of glass coverslips separated by silicone spacers on three sides and filled with RPMI + 10% FBS. For aspiration of the tumour interior, the tumour was sliced with a scalpel or razor blade prior to placement in the chamber. A glass capillary (1 mm outer diameter/0.75 mm inner diameter, World Precision Instruments TW100-3) was pulled on a Browning/Flaming type pipette puller (Sutter Instruments P-97), scored with a ceramic tile, and broken to obtain micropipettes with diameters 40-100 µm. Only cleanly broken tips were used for aspiration. When required, micropipettes were bent with a De Fonbrune microforge. The micropipette was backfilled with 1% (w/v) bovine serum albumin (BSA) in PBS to prevent tissue adhesion and connected to a dual-stage water manometer. Aspiration was applied manually with a syringe (0.5-10 mL) and the pressure difference Δ*P* was measured with a calibrated pressure transducer (Validyne). Time-lapse brightfield microscopy was performed on a TE300 microscope (Nikon) with a 20x/0.5 NA objective and images acquired on an Evolve Delata EMCCD camera (Photometrics) using MicroManager v1.4 software. Tissue elongation *L*(*t*) was determined by image analysis with ImageJ and fit to the standard linear solid model^7^ assuming a wall shape parameter Φ ∼ 2. Tumour strain typically reached a plateau value after several seconds that was maintained for the short timescale of these experiments, allowing the elastic modulus to be approximated as *E* ∼ *ΔP* (*L*/*R_P_*)^-1^ as described previously^8^. Tumouroid flow on longer timescales was modeled by including a viscosity term in the standard linear model^9^.

### Statistical analysis and curve fitting

Statistical analyses and curve fitting were performed in Prism 8.4 (GraphPad) and MATLAB R2020a (MathWorks). Details for each analysis are provided in figure legends, and exact p values are provided where n.s. not significant indicates p > 0.05. Tumouroid and tumour growth data (projected area vs. time) were fit to the exponential growth model (*A* = *A*_0_ e*^kt^* for tumours and *A* = *A*_1_ e*^k^*^(*t*-1)^ for tumouroids) using nonlinear least squares regression with prefactors *A*_0_ or *A*_1_, and *k*, the exponential growth rate. Outliers in samples of the fitted parameter *k* (≤1 per condition, sample sizes ≥24) were identified by ROUT’s method (maximum false discovery rate, Q = 1%) in Prism. Cleaned data were fit to mathematical models described in Extended Data Fig. 3 for the dependence of *k* on macrophage number.

## Data availability

The datasets analyzed during the current study are available in the GEO repository, and further information is available in the cited references in which the datasets were generated.

## Code availability

No custom code central to conclusions of this study was generated.

## Author contributions

Conceptualization: JCA, LJD, DED. Formal Analysis: JCA, LJD. Funding Acquisition: JCA, LJD, BHH, DED. Investigation: JCA, LJD, WZ, BHH, SK, RP, MV. Methodology: JCA, LJD, WZ, SK, RP, BHH, JI, CMA. Resources: DED. Supervision: DED. Visualization: JCA, LJD, DED. Writing: JCA, LJD, DED.

## Acknowledgements

This work was supported by funding from the following sources: NIH R01 HL124106 (DED), U54 CA193417 (DED), NIH F32 CA228285 (LJD), and the NSF GRFP DGE-1845298 (JCA and BHH). The authors thank Dr. Chi Van Dang of the Wistar Institute for providing YUMM2.1 cells and to Dr. Igor Brodsky of Penn Vet for providing editable CIM lines. The authors additionally thank Justine Lee for aiding in development of the gene editing pipeline and Dr. Sunny Shin of Penn Vet for support on CIM progenitors. The authors acknowledge the following University of Pennsylvania core facilities: Cell Center Stockroom, Stem Cell & Xenograft Core, Molecular Pathology & Imaging Core, the Penn Vet Comparative Pathology Core, the Cell & Developmental Biology Microscopy Core, the Flow Cytometry & Cell Sorting Facility, and the Penn Genomic Analysis Core.

## Data availability statement

All data are available within the article (and its supplementary information) or available upon reasonable request from the corresponding author.

## Extended Data Figure Legends

**Extended Data Figure 1:**
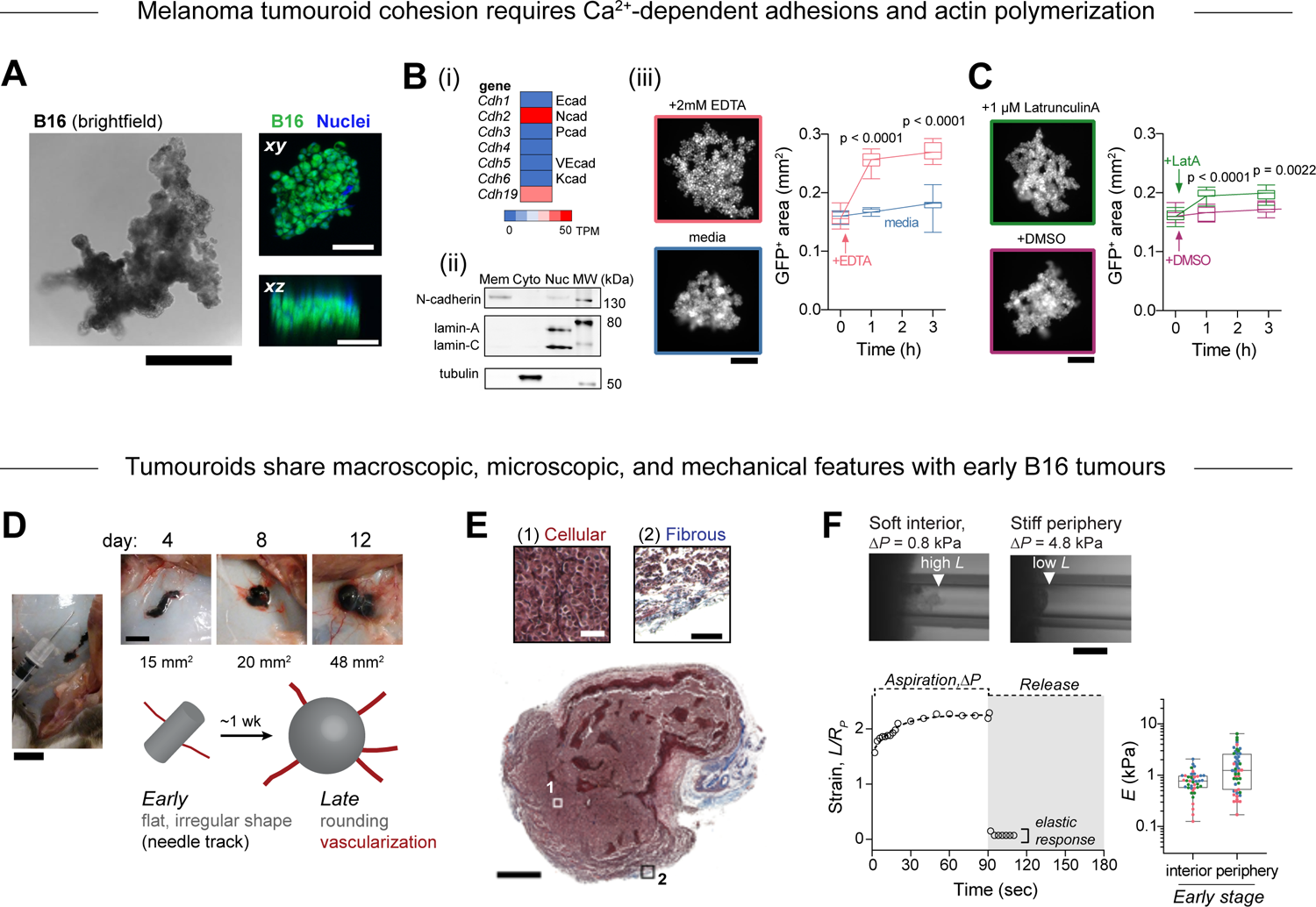
Characterization of B16 tumouroids and subcutaneous tumours. **A** Brightfield and confocal microscopy of B16 tumouroids. (left) Brightfield image of a B16 tumouroid three days after 1000 cells were seeded atop an agarose gel. Large tumouroids typically have irregular borders and appear dark due to melanin pigments. Scale bar: 0.5 mm. (right) Confocal max intensity projections (*xy*, *xz* planes) of a B16 tumouroid GFP (green) and Hoechst nuclear stain (blue). The tumouroid was fixed ∼24 h after 100 cells were seeded in the non-adhesive well. Scale bar: 100 µm. **B** Tumouroid cohesion depends on Ca^2+^. **(i)** Transcriptomic analysis of cultured B16 cells (GSE162105^1^) indicates N-cadherin and cadherin-19 are possible Ca^2+-^dependent cell-cell adhesion molecules. **(ii)** Immunoblotting of B16 lysate fractions confirms N-cadherin protein expression in the membrane fraction and expected lamin-A/C and tubulin distribution in the nuclear and cytoplasmic fractions, respectively. **(iii)** Treatment of B16 tumouroids with 2 mM EDTA results in a rapid (<1 h) increase in tumouroid projected area versus untreated tumouroids (n = 7-8 tumouroids, center line, median; box limits, upper and lower quartiles; whiskers, entire range). Statistical significance was assessed by Welch’s t-test (unpaired, two-tailed). Scale bar: 200 µm. **C** Tumouroid cohesion depends on the actin cytoskeleton. Treatment of B16 tumouroids with 1 µM Latrunculin A results in a rapid (<1 h) increase in tumouroid projected area versus treatment with DMSO vehicle (n = 7-8 tumouroids, center line, median; box limits, upper and lower quartiles; whiskers, entire range). Statistical significance was assessed by Welch’s t-test (unpaired, two-tailed). Scale bar: 200 µm. **D** Photographs of subcutaneous B16 tumours in mice dissected on day 4, 8, or 12 after subcutaneous (s.c.) tumour injection. Day 4 tumours were typically elongated compared to more spherical day 8 and day 12 tumours. Elongation was often observed in the direction in which the needle was inserted during subcutaneous injection of the cell suspension (left). Tumour area calculated in ImageJ is reported beneath each photograph. Scale bars: 10 mm (left) and 5 mm (right). **E** Trichrome-stained B16 tumour section. The tumour interior (region 1, inset scale bar: 50 µm) is cellular (maroon-stained cytoplasm and blacked-stained nuclei) with some necrosis. The tumour periphery contains blue-stained ECM fibers (region 2, inset scale bar: 100 µm) and large, round fat droplets. Scale bar: 1 mm. **F** Interior regions of tumours typically exhibit large deformation *L* at low Δ*P* (top left) while peripheral regions of tumours typically exhibit small *L* and require higher ΔP for measurable deformations (top right). Images depict the extension of tumour into the pipette after 90 s of aspiration at the indicated Δ*P*. Scale bar: 50 µm. Tissue strain (*L*(*t*)/*R_P_*) versus time is plotted for the constant aspiration pressure creep phase (0-90 s) and after the pressure was released back to Δ*P* = 0 for a representative interior region of a tumour. The profile is typical of a viscoelastic solid and the deformation was recoverable when the pressure was released. The black line depicts nonlinear regression of *L*(*t*)/*R_P_* to the standard linear solid model, which includes the elastic modulus *E* (as *t* → ∞) as a fitted parameter. Elastic moduli (*E*) from interior and peripheral regions of an early-stage (day 4) tumour were calculated from the plateau in the strain vs. time plot. Each symbol represents a separate aspiration-release measurement. The symbol colors correspond to three different tumours (15-30 measurements per tumour, center line, median; box limits, upper and lower quartiles; whiskers, entire range).

**Extended Data Figure 2:**
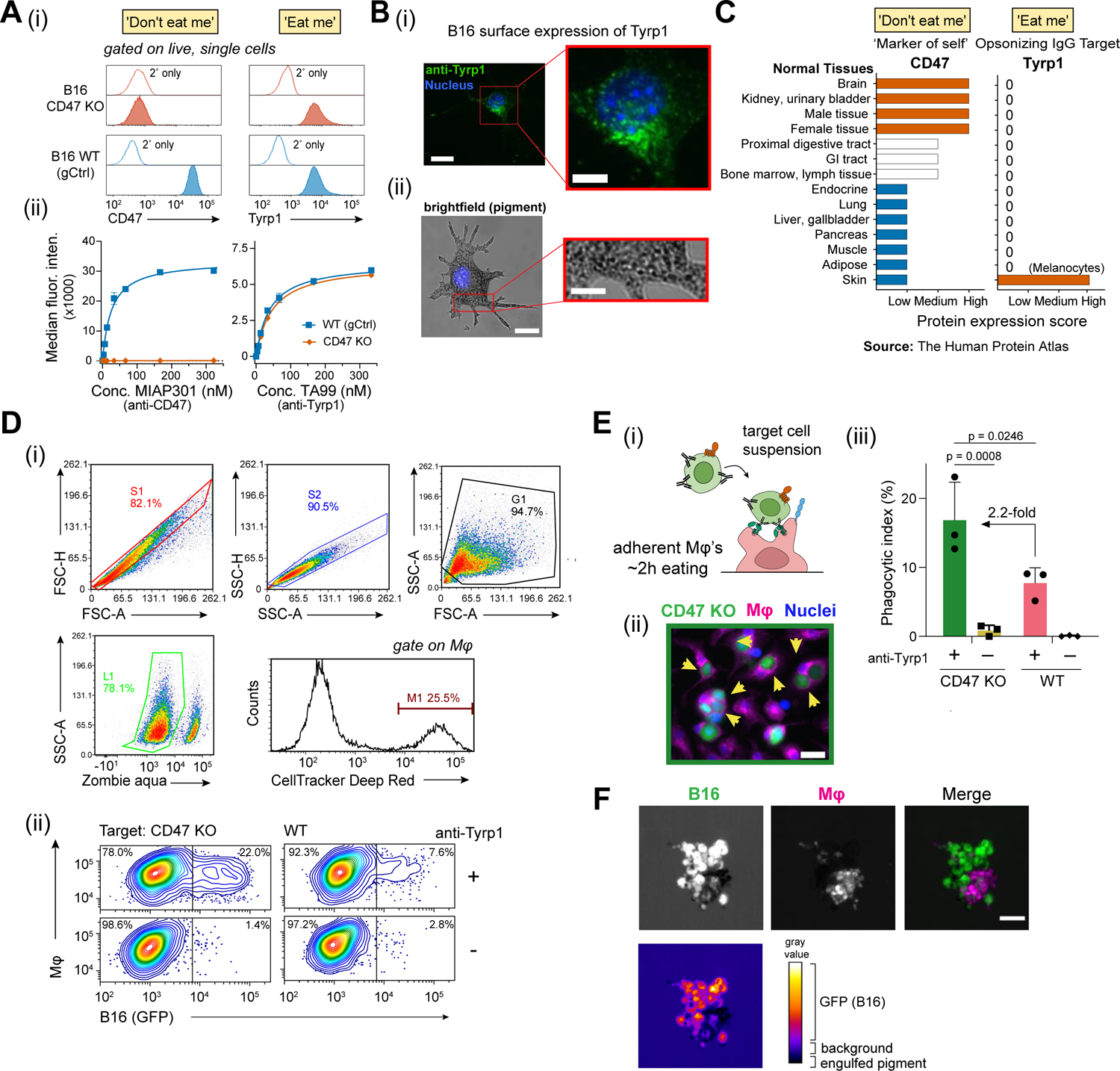
Analysis of CD47 and Tyrp1 expression and phagocytosis of B16 tumouroids and cell suspensions. **A** Flow cytometry analysis of CD47 and Tyrp1 expression on B16 cell lines. **(i)** Representative histograms of anti-CD47 (left) and anti-Tyrp1 (right) binding to CD47 KO (top) and WT (bottom) B16 cell lines, which were detected with secondary antibodies conjugated with Alexa Fluor 647. The populations shown in the histograms were gated on live, single cells using forward/side scatter and DAPI staining. **(ii)** Median fluorescence intensity calculated for CD47 KO and WT cells incubated with varying concentrations of anti-CD47 or anti-Tyrp1 followed by secondary antibody (n = 3 tests per antibody concentration, mean ± SD). The data were fit to a hyperbolic binding model of the form *y* = *A*\**x*/(*K*+*x*), which is shown as a solid line. **B** Tyrp1 protein expression on B16 melanoma cells. **(i)** Immunofluorescence image of a B16 cell surface-stained (fixed, unpermeabilized) with anti-Tyrp1 and anti-mouse IgG Alexa Fluor 647 (green) and counterstained with Hoechst 33342 (blue). **(ii)** Brightfield image of a B16 cell showing pigmented melanosomes. Scale bars: 25 µm (left) and 10 µm (right). **C** Protein expression data from The Human Protein Atlas for CD47 (left) and Tyrp1 (right) across different normal tissues. CD47 is expressed in all tissues consistent with its role as a ‘marker of self’. Tyrp1 expression is limited to skin, and specifically to melanocytes. **D** Representative flow cytometry analysis of disaggregated tumouroids to measure phagocytosis. **(i)** Flow cytometry events were gated on live, single cells using forward/side scatter and Zombie aqua viability dye. Macrophages were gated based on CellTracker Deep Red fluorescence. **(ii)** The % phagocytic macrophages reported in Fig. 1C is calculated as the number of DeepRed^+^GFP^+^ double positive events divided by all DeepRed^+^ events multiplied by 100. **E (i)** Conventional 2D phagocytosis of WT and CD47 KO B16 cell lines by primary mouse bone marrow-derived macrophages (BMDMs). **(ii)** B16 cells (green) were incubated with opsonizing anti-Tyrp1 or IgG2a isotype control antibody and added to adherent BMDMs (magenta). Yellow arrows denote phagocytic events, with some BMDMs engulfing multiple KO cells. Scale bar: 25 µm. **(iii)** The phagocytic index is calculated as the percentage of macrophages engulfing a target cell multiplied by the number of engulfed cells per macrophage (mean ± SD, n = 3 wells). Statistical significance was assessed by two-way ANOVA and Tukey’s multiple comparison test. **F** Phagocytic macrophages in tumouroids co-localize with melanin pigments, which is most easily visualized as a dark shadow in the GFP channel. The signal intensity in this region is below background light intensity, consistent with light absorbance by pigments. Scale bar: 100 µm.

**Extended Data Figure 3:**
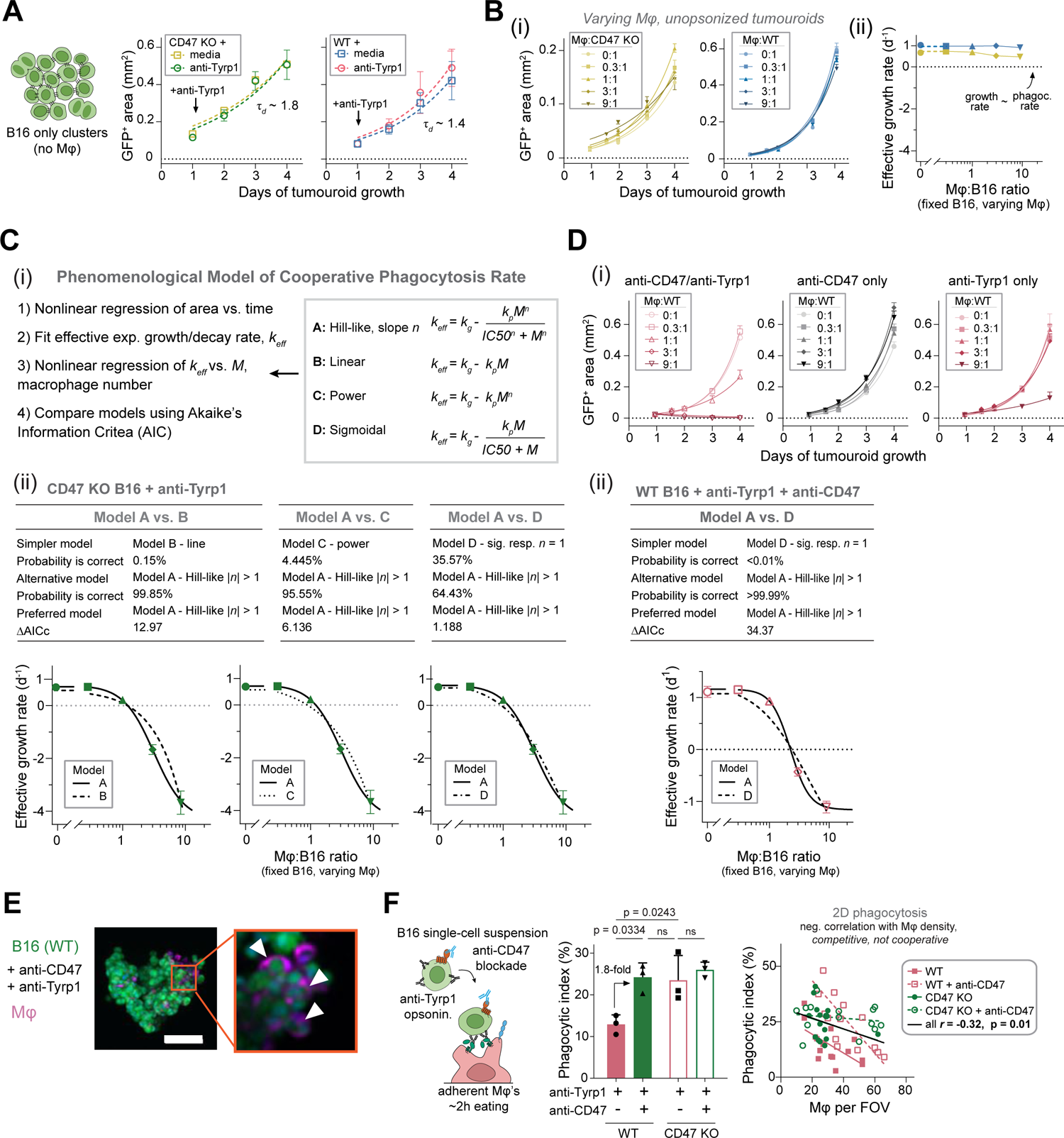
Modeling cooperative tumouroid phagocytosis and non-cooperative phagocytosis in the conventional 2D assay. **A** Tumouroid growth is unaffected by anti-Tyrp1 added on day 1 without macrophages, indicating that anti-Tyrp1 has no direct cytotoxicity and that macrophages are required to mediate its effects (mean ± SD, n = 6 tumouroids). **B (i)** Macrophages do not significantly repress or eliminate CD47 KO or WT tumouroids in the absence of anti-Tyrp1 at any macrophage:B16 ratio tested (mean ± SEM, n = 6-8 tumouroids in a representative experiment). **(ii)** The effective growth is greater than 0 and shows little dependence on the macrophage:B16 ratio for unopsonized tumouroids (mean ± SEM, n ≥ 24 tumouroids across the four independent experiments shown in Fig 1E). **C** Comparison of mathematical models for phagocytosis of tumouroids. **(i)** Workflow for fitting *k_eff_* from tumouroid area (*A*) growth or decay curves (Fig 1E) and subsequent model fitting of *k_eff_* vs. macrophage number (*M*). Four models (models A-D) were considered including (model A) a Hill-like sigmoidal response on *M*, (model B) linear dependence on *M*, (model C) a power law dependence on *M*, and (model D) a sigmoidal dependence on the number of *M*. **(ii)** The fits of models B-D were compared one-by-one to model A using Akaike’s Information Criteria (AIC) to determine whether the four-parameter Hill-like model A is justified rather than comparatively simpler models with two or three parameters. The comparisons are summarized in the tables and the model fits to *k_eff_* vs. macrophage:B16 ratio data (Fig. 1E) are plotted beneath. **D (i)** Growth curves for WT tumouroids treated with anti-CD47 and/or anti-Tyrp1 that correspond to the plot of *k_eff_* vs macrophage:B16 ratio in Fig 1F-i. **(ii)** The fits of models A and D were compared using Akaike’s Information Criteria (AIC). The comparison is summarized in the tables and the fits are plotted beneath. **E** Representative max intensity projection of confocal images of a WT tumouroid one day after addition of macrophages, anti-Tyrp1, and anti-CD47. Scale bar 100 µm. The expansion depicts multiple macrophages that have engulfed B16 (denoted by white arrowheads). **F** Conventional 2D phagocytosis assay with adherent macrophages and single-cell suspensions of WT or CD47 KO B16 cells opsonized with anti-Tyrp1 and treated with either anti-CD47 or rIgG2a isotype control antibody (mean ± SD, n = 3 wells per condition). Statistical significance was assessed by two-way ANOVA and Tukey’s multiple comparison test. Across all four conditions (2 cell lines x 2 blocking conditions), the number of macrophages per field of view (FOV) is plotted against the phagocytic index calculated for that FOV, revealing a weak negative correlation (Pearson’s *r* = −0.32, p = 0.01).

**Extended Data Figure 4:**
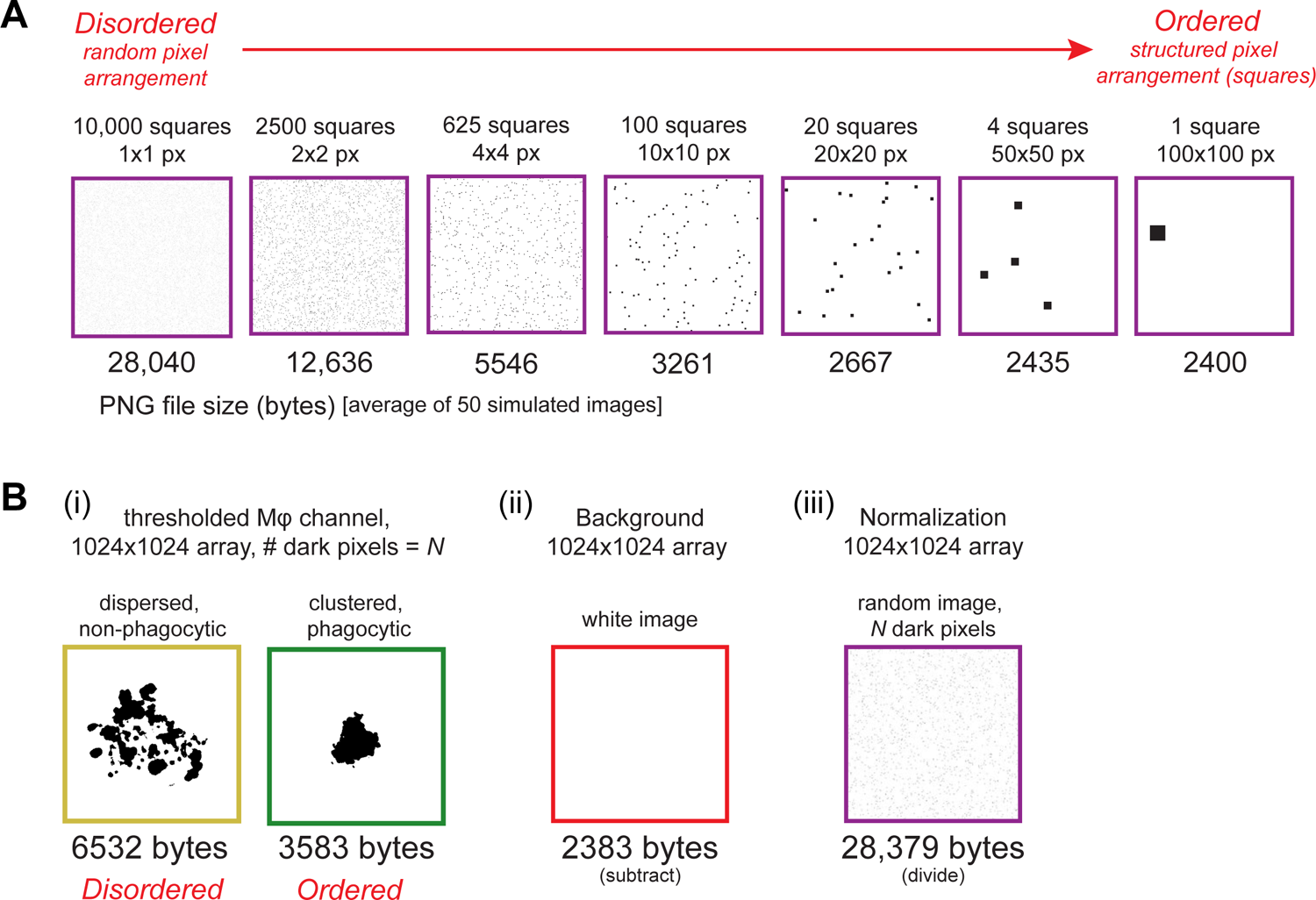
Informational entropy calculation of macrophage order based on image compression. **A** Simulated 1024×1024 pixel (px) images containing 10,000 dark pixels on a white background that are arranged in random squares varying in size from 10,000 1×1 px squares to one 100×100 px square. The images were compressed in the portable network graphics (PNG) file format. The PNG file size (in bytes) below each image is the average of 50 randomly generated images with the same specifications. File size decreases with increasing image order/entropy. **B** To compare images with different numbers of dark pixels (*N*), a normalization procedure involved subtracting the PNG file size of a completely white image of the same dimension and then dividing this value by the PNG file size of an equal dimension random image with *N* dark pixels.

**Extended Data Figure 5:**
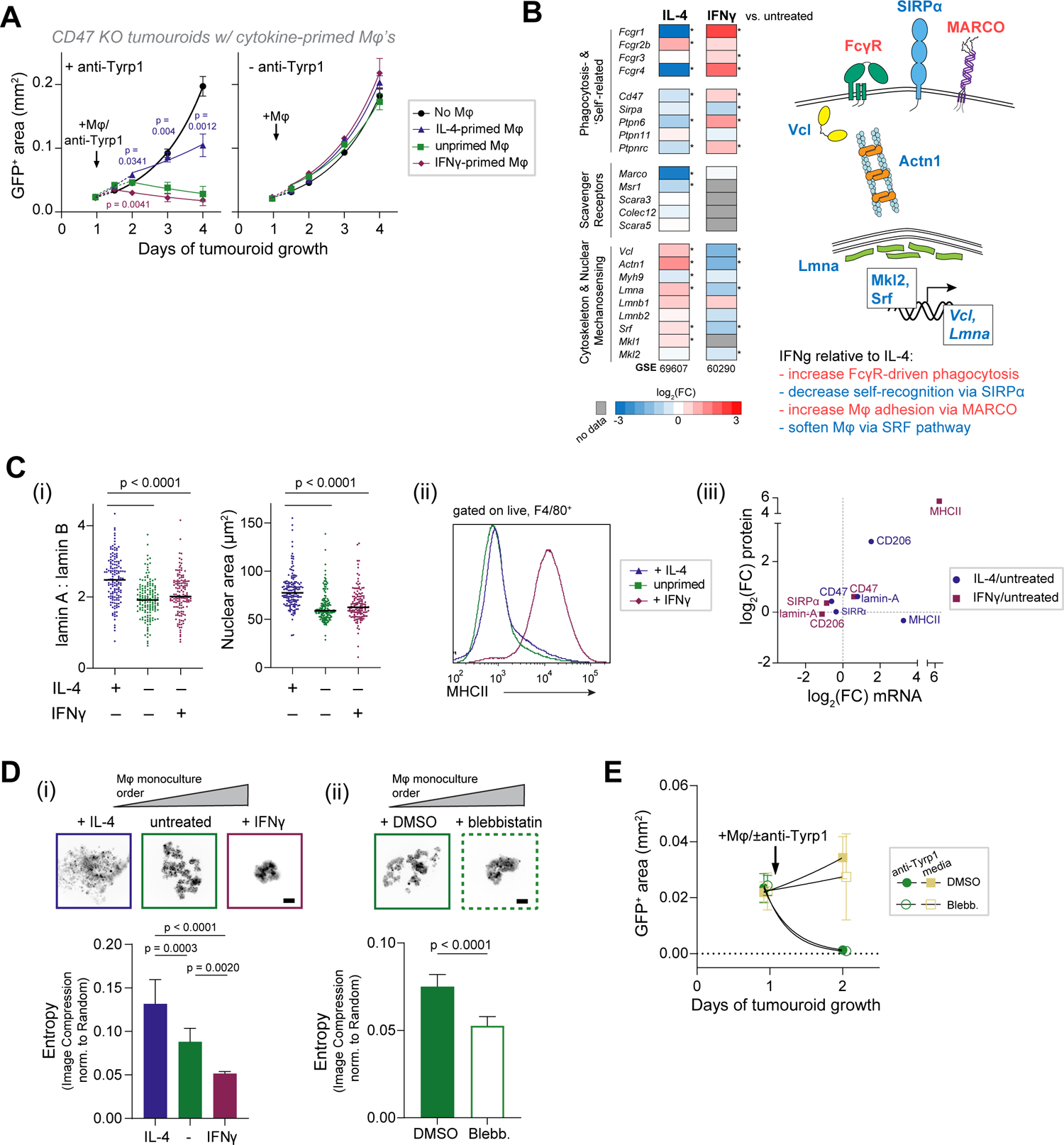
Effects of cytokine priming or the myosin-II inhibitor blebbistatin on tumouroid growth suppression and macrophage clustering. **A** Tumouroid growth or repression following addition of macrophages primed with either 20 ng/mL IFNγ or 20 ng/mL IL-4 for 48 h with (left) or without (right) anti-Tyrp1. (mean ± SEM, n = 7-8 tumouroids). Statistical significance for each time point was assessed by one-way ANOVA and Dunnett’s multiple comparison test between unprimed and IFNγ-primed or IL-4-primed macrophages. **B** Transcriptomic microarray analyses of BMDMs treated with IFNγ or IL-4 versus untreated BMDMs (GSE60290^2^, GSE69607^3^). The heat map depicts log_2_(fold change) for selected genes. Proteins encoded by differentially expressed genes are depicted in the schematic detailing a putative cytoskeleton and nuclear mechanosensing pathway and membrane receptors involved in phagocytosis and macrophage adhesion. Statistical significance was assessed by computing an adjusted p-value by the method of Benjamini and Hochberg (* p < 0.05). **C** Measurements of protein expression of selected differentially expressed genes in BMDMs treated with IFNγ or IL-4 for 48 h. **(i)** The ratio of lamin-A to lamin-B and the projected nuclear area were quantified by immunofluorescence microscopy (n > 140 cells per condition across 8 fields of view). Statistical significance was assessed by one-way ANOVA followed by Tukey’s multiple comparison test. **(ii)** Surface expression of MHCII in IFNγ- or IL-4-treated or untreated BMDMs was measured by flow cytometry. Similar staining was also performed with antibodies against CD206, CD47, and SIRPα. **(iii)** The fold change (cytokine-primed vs. untreated) of indicated proteins from immunofluorescence microscopy and flow cytometry experiments is plotted against the fold change in gene expression from microarrays. **D** Entropy analysis for **(i)** cytokine-primed and **(ii)** blebbistatin-treated macrophages cultured on non-adhesive surfaces at a density of 1000 cells per well (mean ± SD, n = 8 wells). Statistical significance was assessed by one-way ANOVA followed by Tukey’s multiple comparison test for cytokine priming and Welch’s t-test for blebbistatin treatment. Cytokines (20 ng/mL) or blebbistatin (20 µM) were added after 6 h and images were acquired after 48 h. Scale bars: 100 µm. **E** Tumouroid growth repression by macrophages was not affected by addition of 20 µM blebbistatin (mean ± SD, n = 3-4 tumouroids).

**Extended Data Figure 6:**
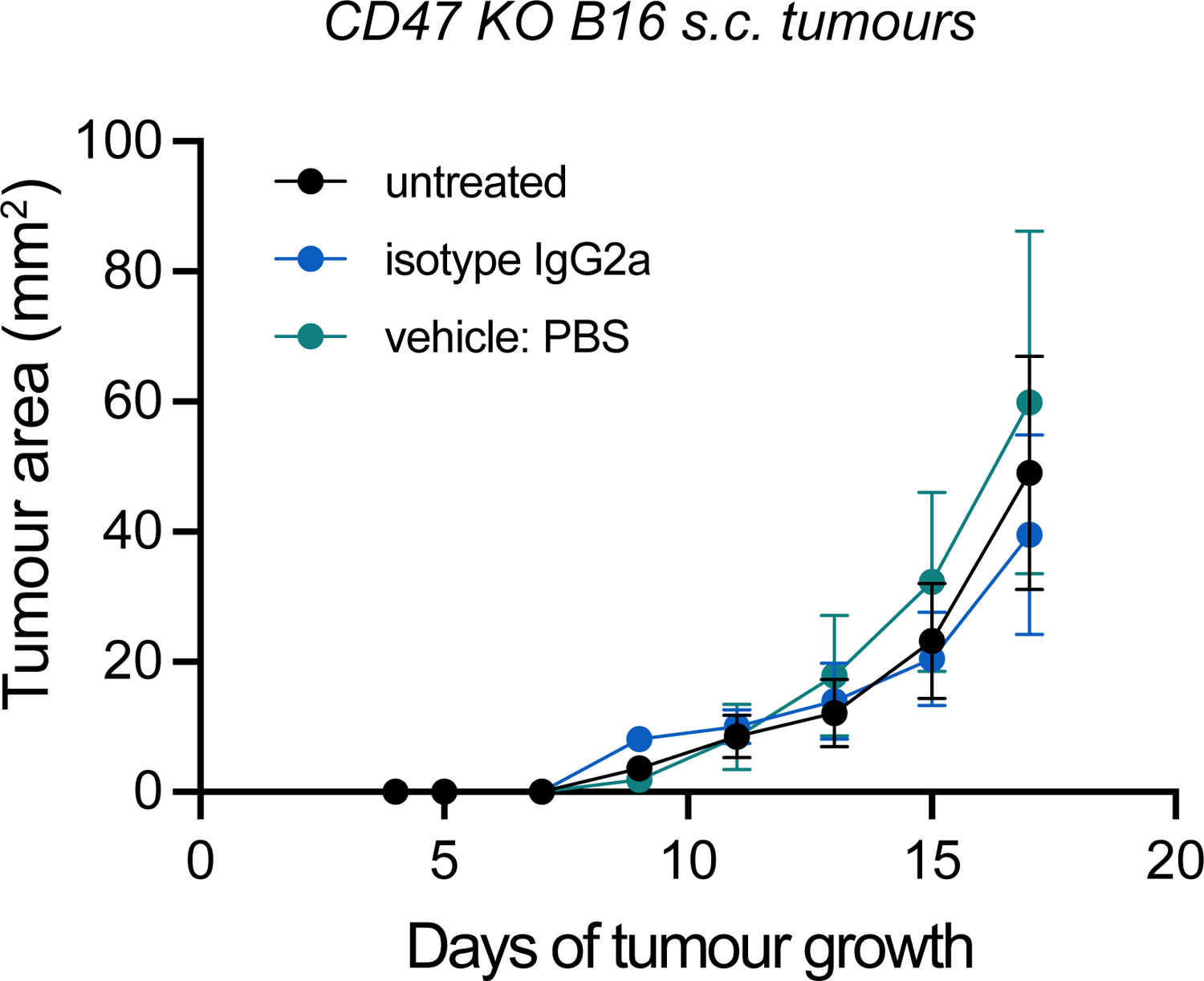
Isotype IgG2a control or vehicle injections have no effect on CD47 KO tumour growth relative to untreated tumours. Growth curves of mice bearing CD47 KO B16 tumours (2×10^5^ s.c. flank injection) and treated i.v. on days 4, 5, 7, 9, 11, 13, and 15 with 250 µg isotype IgG2a control (clone C1.18.4), 100 µl PBS vehicle, or left untreated. Clone C1.18.4 is the isotype control antibody for anti-Tyrp1 clone TA99. There are no significant growth differences (mean ± SEM, n = 6 mice per group).

**Extended Data Figure 7:**
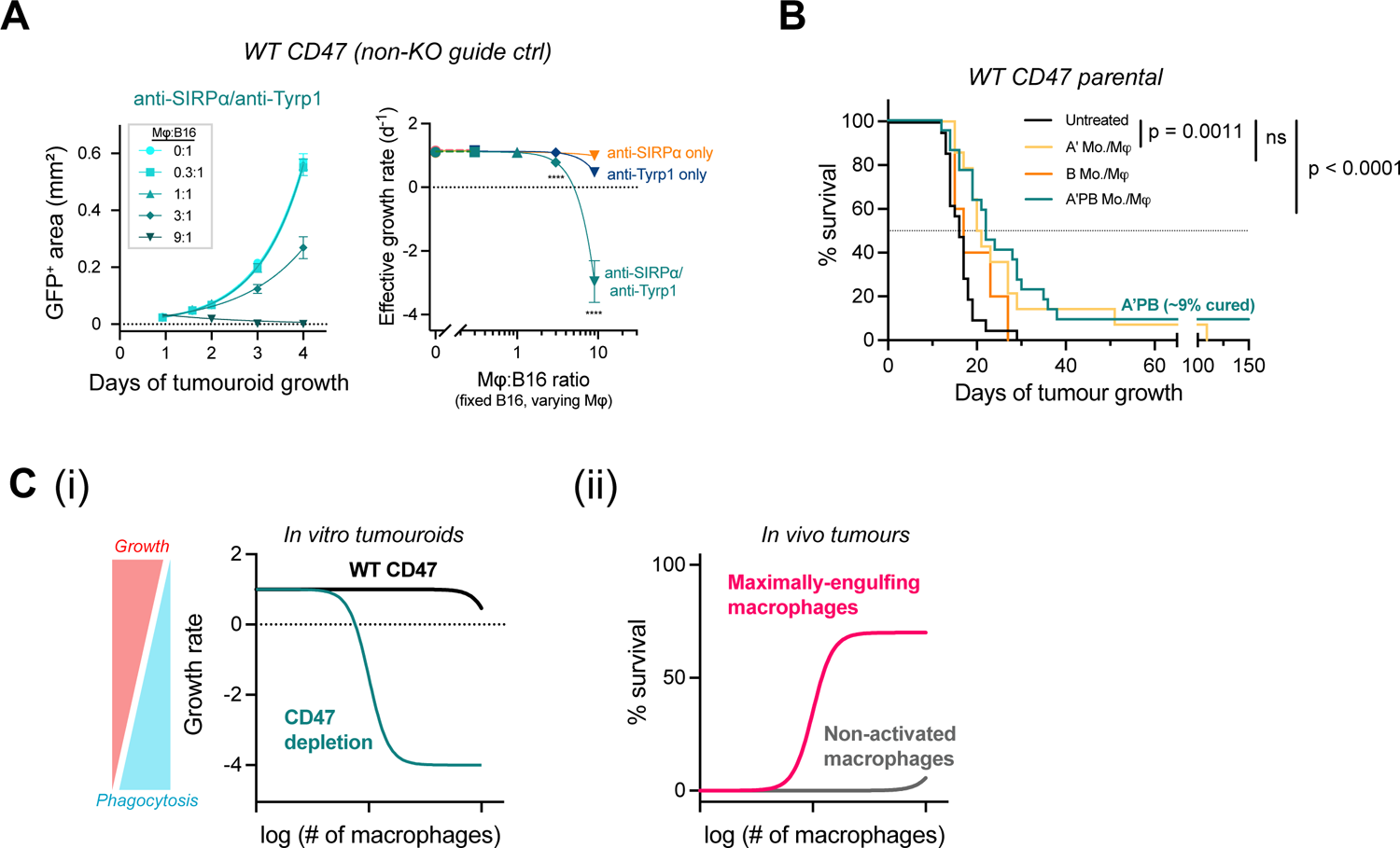
Fc Receptor-primed, SIRPα-blocked (A’PB) marrow cells elicit complete anti-tumour responses in a fraction of mice bearing WT B16 tumours, while control marrow cells cannot. **A** A’PB macrophages in tumouroids can effectively repress growth at high macrophage:B16 (WT CD47 – non-KO guide ctrl) ratios, while anti-SIRPα (B) and anti-Tyrp1 (A’) only conditions fail to control tumouroids. **B** Survival curves of mice bearing WT B16-F10 tumours (parental line) and treated with 2×10^5^ A’PB, A’ (Fc-receptor primed only, no SIRPα block), or B (no Fc-receptor priming, only SIRPα block) marrow cells i.v. on day 4 after tumour engraftment relative to untreated mice. A’PB and A’ conditions received additional 250 µg anti-Tyrp1 i.v. on days 5, 7, 9, 11, 13, and 15. **C (i)** Summary of macrophage number versus therapeutic outcome for tumouroids *in vitro* and tumours *in vivo*. Growth rate is controlled by tumour cell proliferation and macrophage phagocytosis. **(ii)** CD47 signaling dominates pro-phagocytic signaling at low and high macrophage numbers, but CD47 depletion, IgG opsonization, and macrophage infiltration can eliminate tumours.

**Extended Data Figure 8:**
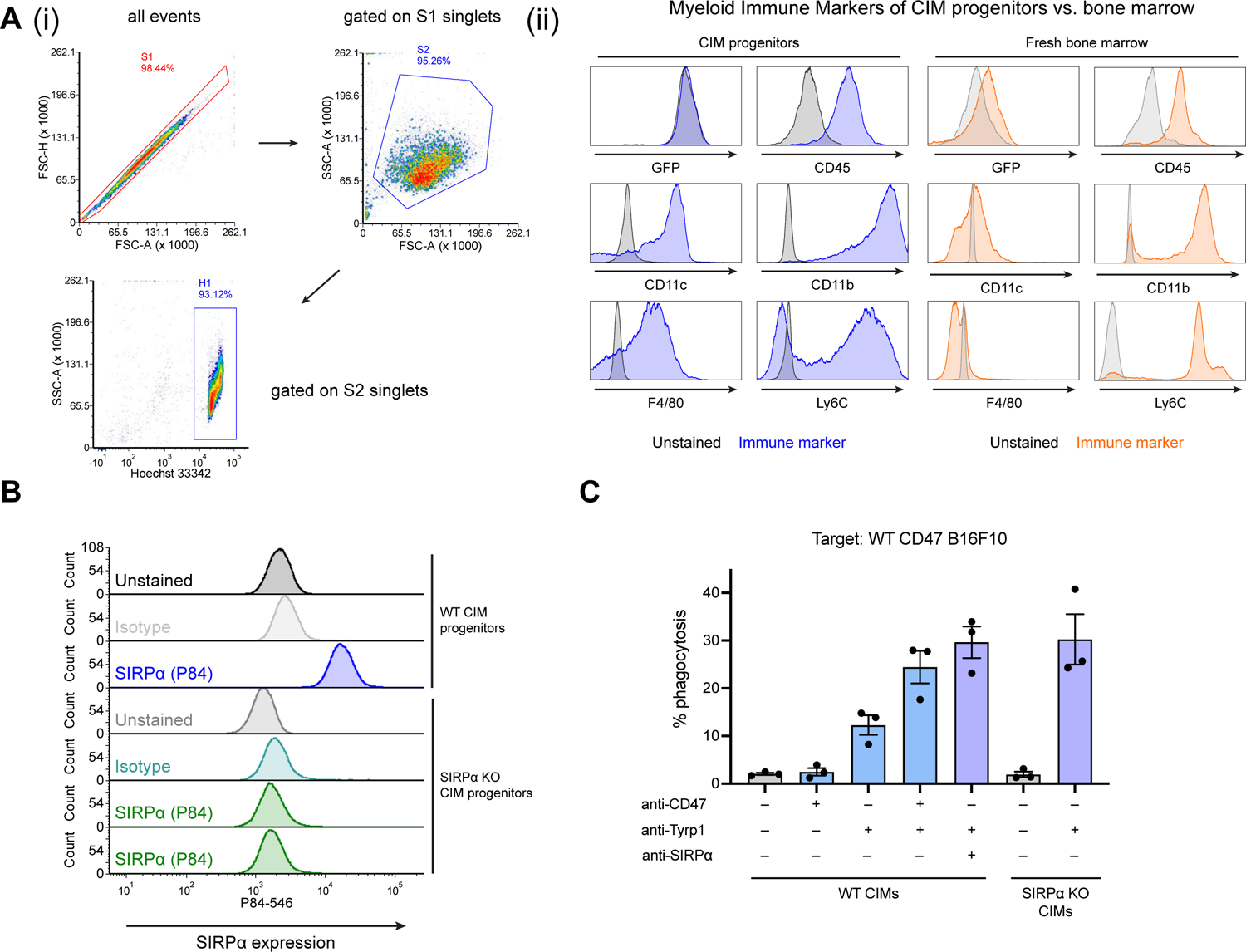
Conditional immortalized macrophage (CIM) progenitors are myeloid committed lineage cells that can be differentiated to phagocytic macrophage phenotype. **A (i)** Gating strategy for flow cytometry evaluation of CIM progenitors. **(ii)** CIM progenitors express myeloid-specific immune markers. Progenitors are GFP+ from their source marrow from the Rosa26-Cas9 knock-in mouse on the C57BL6/J background.^4,5^ Immune marker staining on freshly harvested bone marrow cells is shown for comparison with lower F4/80 expression due to less lineage commitment compared to CIM progenitors. **B** Flow histograms showing that SIRPα KO CIM progenitors are depleted of SIRPα relative to unedited wild-type CIM progenitors.

**Extended Data Figure 9:**
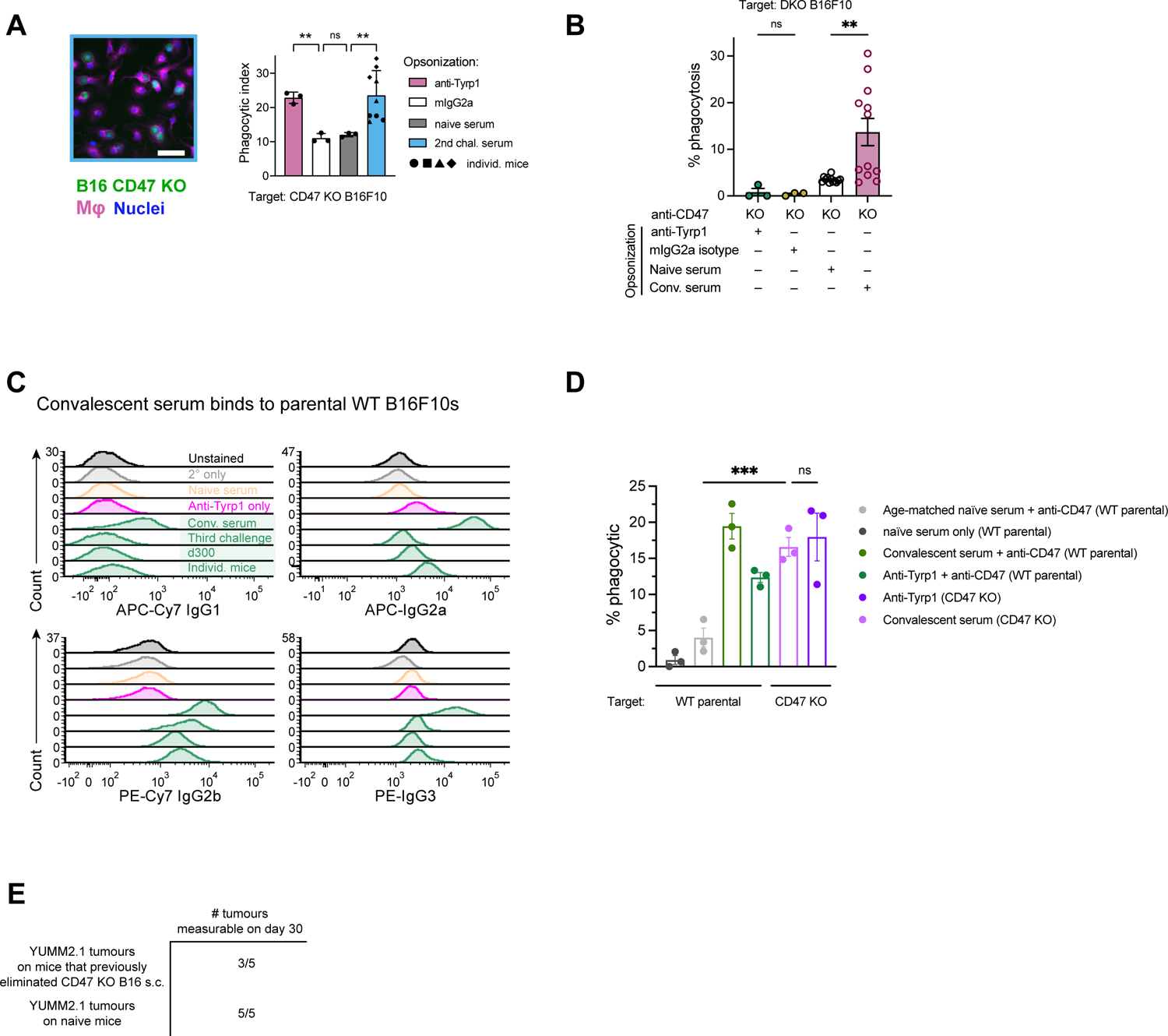
Convalescent serum IgG from CD47 KO tumour complete responders binds and opsonizes both CD47 KO and WT CD47 B16s. **A** Representative fluorescent image (left) showing serum IgG-opsonized CD47 KO B16s engulfed by BMDMs. Convalescent serum IgG opsonizes CD47 KO B16s for phagocytosis by BMDMs as effectively or more highly than anti-Tyrp1 (right). Each serum symbol shape represents a different mouse sample (n = 3 per mouse sample, mean ± SEM). Statistical significance was assessed by Brown-Forsythe and Welch ANOVA tests with Dunnett’s T3 multiple comparisons test between selected groups. **B** Convalescent serum IgG from third challenge complete responder mice opsonizes CD47 KO B16s and DKO B16s that lack Tyrp1 by BMDMs in 2D phagocytosis assays (same mouse serum samples from 3D assays in Figure 4D). Statistical significance was assessed by one-way ANOVA and Sidak’s multiple comparison test between selected groups, ns not significant (n = 3 for antibody conditions, n = 12 for sera conditions, mean ± SEM). **C** Flow histograms of 4 (CD47 KO) third challenged (serum drawn 300 days after tumour challenge) mice showing that IgG in convalescent serum from complete responder mice binds to WT B16s. Anti-B16 IgG2a and IgG2b subclasses bind equivalently to or more highly than anti-Tyrp1. **D** Serum IgG derived from CD47 KO tumour complete responders retains opsonization and functional phagocytic ability against WT B16s. Statistical significance was assessed by one-way ANOVA and Tukey’s multiple comparison test between selected groups, ns not significant (n = 3 per condition, mean ± SEM). **E** Mice that showed complete anti-tumour responses after three challenges of CD47 KO B16 tumours also grow YUMM2.1 tumours.

**Extended Data Figure 10:**
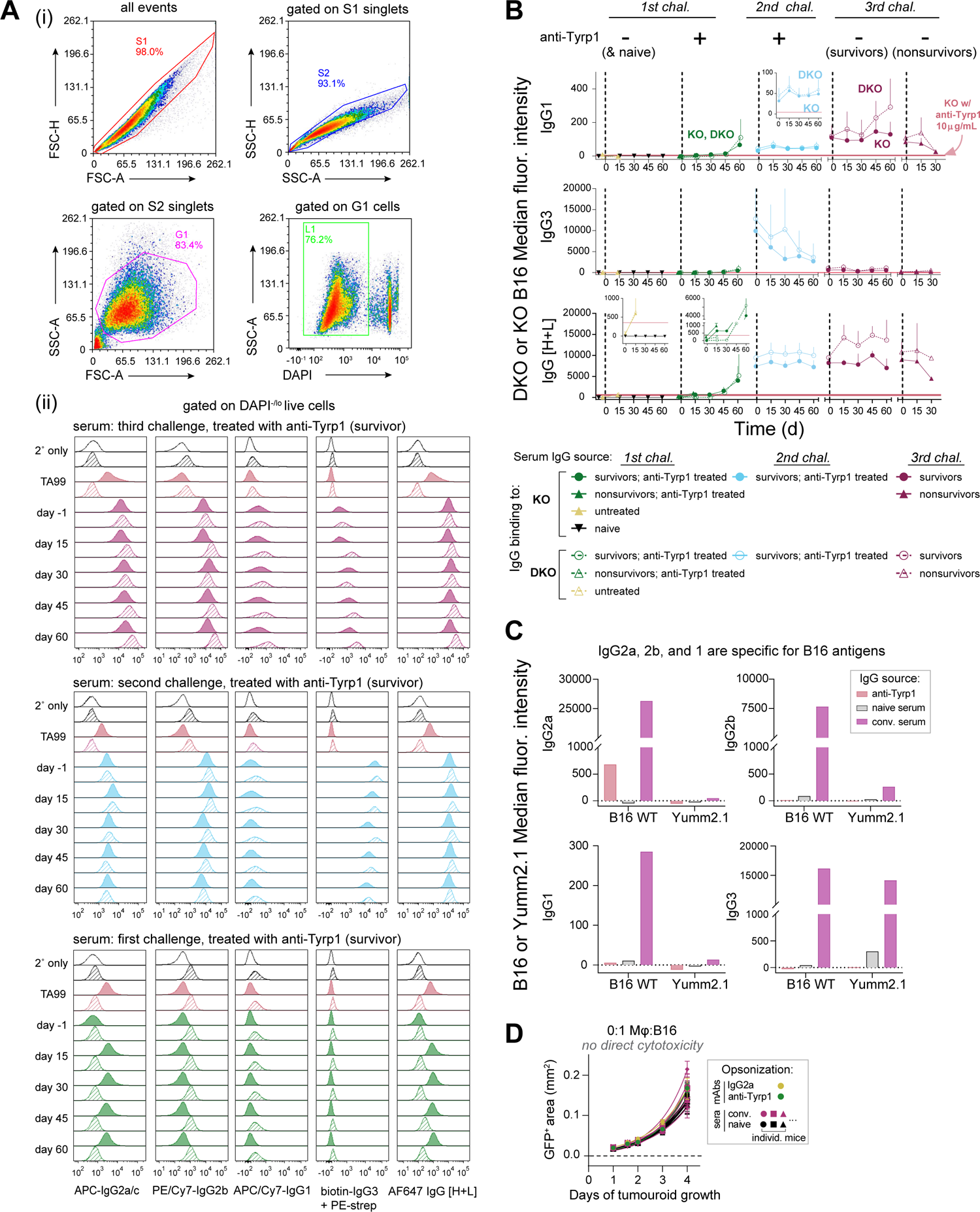
Convalescent serum IgG from CD47 KO tumour complete responders shows subclass-specific binding to CD47 KO B16s and DKO B16s that lack Tyrp1, and no binding of IgG1, IgG2a/c, and IgG2b to non-B16 YUMM2.1 cells. **A** Flow cytometry analysis of serum IgG binding to B16 cell lines. **(i)** General gating strategy for flow cytometry analysis of IgG binding to B16 cell lines. Doublet discrimination was performed with pulse height and pulse area parameters in the forward and side scatter channels (S1 gate, FSC-H vs. FSC-A) and (S2 gate, SSC-H vs. SSC-A). Cells were distinguished from debris as shown in the G1 gate in the SSC-A vs. FSC-A plot). Dead cells were excluded from analyses by DAPI staining when the analysis was to be performed immediately on viable cells or by Zombie aqua dye when the analysis was to be performed at a later time after fixing cells. **(ii)** Representative flow cytometry histograms of KO (solid filled) and DKO (diagonal hashed) cells incubated with 5% (v/v) serum collected from mice undergoing KO tumour 1^st^, 2^nd^, or 3^rd^ challenge on days −1, 15, 30, 45, and 60 after tumour inoculation. Cells were stained with a panel of monoclonal secondary antibodies specific for mouse IgG1, IgG2a/c, IgG2b, and IgG3 or with a polyclonal secondary antibody recognizing heavy and light chains [H+L] of all mouse IgG subclasses. Control samples were stained with either secondary antibodies only or 10 µg/mL anti-Tyrp1. Binding of anti-Tyrp1 was detected only with the IgG2a/c and polyclonal IgG [H+L] secondary antibodies. **B** Kinetics of *de novo* IgG generation revealed by flow cytometry analysis of KO (filled symbols) and DKO cells (open symbols) stained as described in **A-ii** with serum and with secondary antibodies specific for mouse IgG1 and IgG3 (rows 1-2) or a polyclonal anti-mouse IgG [H+L] secondary antibody (row 3). The corresponding data for IgG2a/c and IgG2b are shown in Fig. 4B. The median fluorescence intensity was corrected by subtracting the background fluorescence equal to the signal of cells incubated with secondary antibodies only. Symbols and error bars depict the mean ± SEM of each group. The range of staining observed with 10 µg/mL anti-Tyrp1 is denoted by the pink boxes or lines. **C** Median fluorescence intensity of B16 WT or Yumm2.1 cells incubated with 5% (v/v) serum collected from a mouse that survived the third challenge, with naïve serum, or with anti-Tyrp1 followed by staining with a panel of monoclonal secondary antibodies specific for mouse IgG1, IgG2a/c, IgG2b, and IgG3. **D** Convalescent sera do not exhibit direct cytotoxic toward KO tumouroids and require macrophages to suppress growth and eliminate tumouroids (Fig. 4D). Tumouroids treated with convalescent serum, naïve serum, anti-Tyrp1, and IgG2a isotype control all grow similarly. Solid lines are nonlinear regression of the data to a simple exponential of the form *A*(*t*) = *A*_1_ exp[*k_eff_*(*t*-1)] (mean ± SEM, n = 3 or 4 tumouroids per sample for 9 convalescent and 9 naïve sera).

**Extended Data Figure 11:**
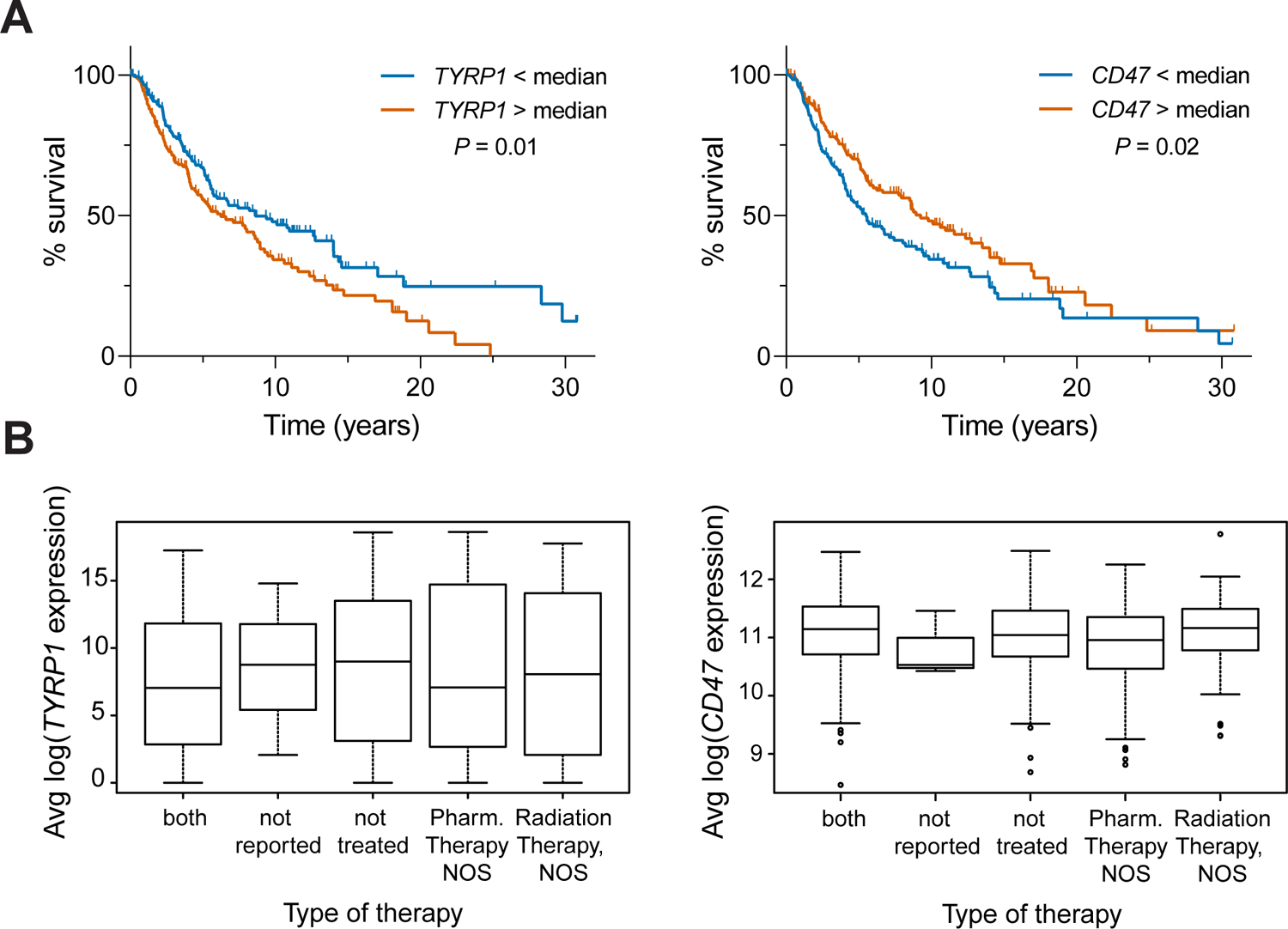
TCGA analysis of TYRP1 and CD47 in human melanoma. **A** Survival analysis of metastatic melanoma patients in The Cancer Genome Atlas (TCGA) based on expression of *TYRP1* (left) and CD47 (right) above or below the median. Significance was determined by the Log-rank (Mantel-Cox) test. **B** Expression levels of *TYRP1* (left, p = 0.68, one-way ANOVA) and *CD47* (right, p = 0.36) were not dependent on treatment reported in TCGA.

**Extended Data Figure 12:**
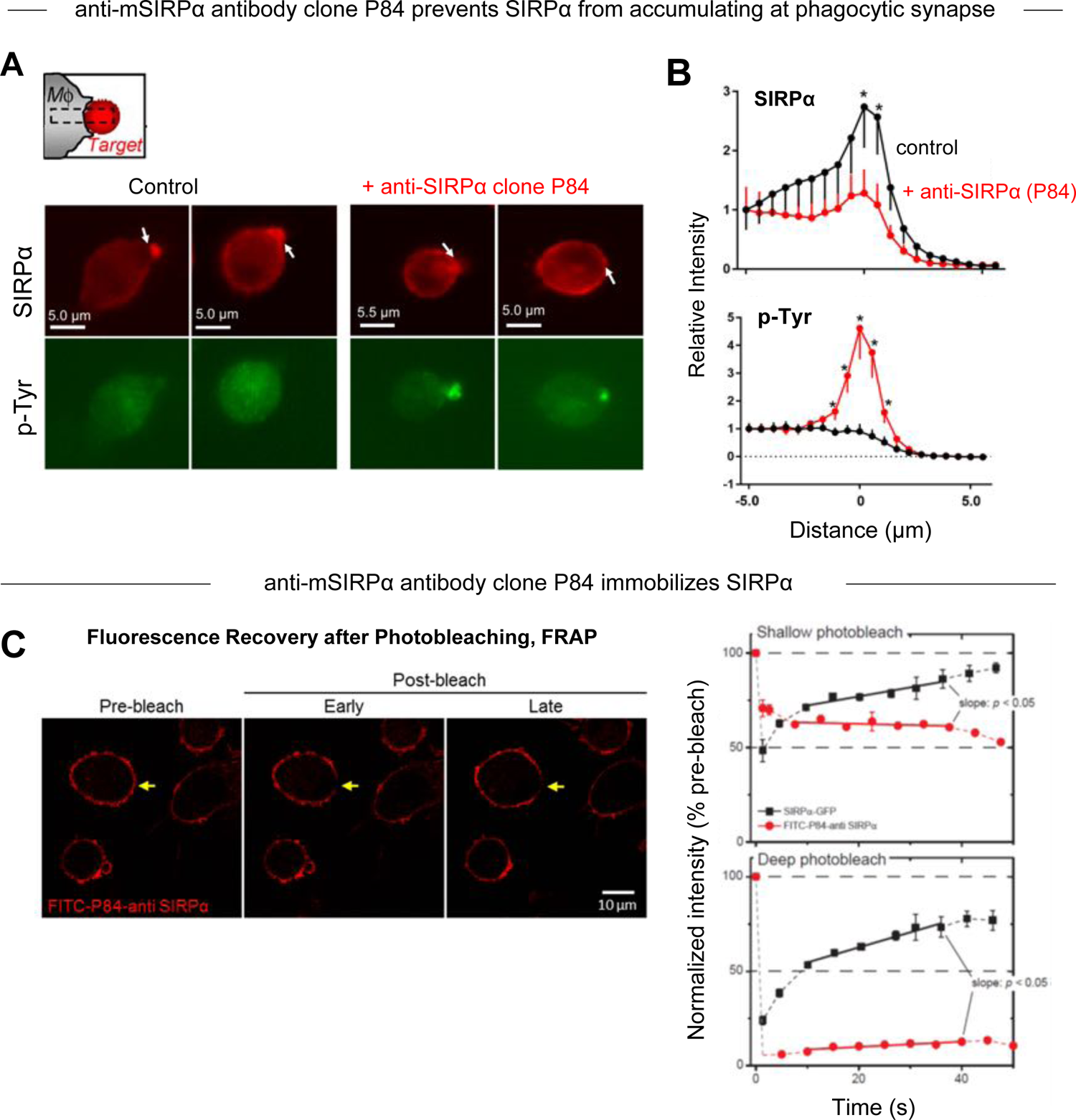
Anti-mouse SIRPα antibody clone P84 binds to mouse macrophages and immobilizes SIRPα. **A** Representative images of mouse macrophages ± anti-mSIRPα (P84) incubated with opsonized mouse RBCs for 30 minutes. Cells were fixed and stained for SIRPα and phospho-tyrosine. **B** Quantification of SIRPα and phospho-tyrosine accumulation at the phagocytic synapse of mouse macrophages ± anti-mSIRPα clone P84. Fluorescence intensity was quantified in each phagocytic synapse as previously described^6^ normalized to macrophages that were not undergoing phagocytosis (n = 7). **C** Representative fluorescence images and FRAP analysis SIRPα-GFP diffusion ± anti-mSIRPα clone P84. Sides of cells were bleached and allowed to recover (n = 3).

